# Lymphatic muscle cells are unique cells that undergo aging induced changes

**DOI:** 10.1101/2023.11.18.567621

**Authors:** Pin-Ji Lei, Katarina J. Ruscic, Kangsan Roh, Johanna J. Rajotte, Meghan J. O’Melia, Echoe M. Bouta, Marla Marquez, Ethel R. Pereira, Ashwin S. Kumar, Guillermo Arroyo-Ataz, Mohammad S. Razavi, Hengbo Zhou, Lutz Menzel, Heena Kumra, Mark Duquette, Peigen Huang, James W. Baish, Lance L. Munn, Jessalyn M. Ubellacker, Dennis Jones, Timothy P. Padera

## Abstract

Lymphatic muscle cells (LMCs) within the wall of collecting lymphatic vessels exhibit tonic and autonomous phasic contractions, which drive active lymph transport to maintain tissue-fluid homeostasis and support immune surveillance. Damage to LMCs disrupts lymphatic function and is related to various diseases. Despite their importance, knowledge of the transcriptional signatures in LMCs and how they relate to lymphatic function in normal and disease contexts is largely missing. We have generated a comprehensive transcriptional single-cell atlas—including LMCs—of collecting lymphatic vessels in mouse dermis at various ages. We identified genes that distinguish LMCs from other types of muscle cells, characterized the phenotypical and transcriptomic changes in LMCs in aged vessels, and uncovered a pro-inflammatory microenvironment that suppresses the contractile apparatus in advanced-aged LMCs. Our findings provide a valuable resource to accelerate future research for the identification of potential drug targets on LMCs to preserve lymphatic vessel function as well as supporting studies to identify genetic causes of primary lymphedema currently with unknown molecular explanation.

## Introduction

The lymphatic vascular system plays a crucial role in maintaining tissue-fluid balance, absorbing dietary lipids, and transporting antigens and immune cells to lymph nodes (Petrova and Koh, 2018). Lymph—which is composed of absorbed interstitial fluid, macromolecules and immune cells—is formed by initial lymphatic vessels and drained into collecting lymphatic vessels, which transport the lymph to lymph nodes and eventually back to the bloodstream (Alitalo, 2011; Mäkinen et al., 2021; Oliver et al., 2020; Petrova and Koh, 2018, 2020). Initial lymphatic vessels are composed of a single layer of oak-leaf shaped lymphatic endothelial cells (LECs) sitting on a fenestrated basement membrane with button-like junctions between LECs (Baluk et al., 2007; Breslin et al., 2019). These button-like junctions allow for the collection of interstitial fluid and macromolecules to form lymph (Trzewik et al., 2001). Muscle cells are absent in the wall of initial lymphatic vessels (Baluk et al., 2007; Mäkinen et al., 2021). In contrast, the wall of collecting lymphatic vessels has three layers: an inner layer of endothelium, a middle layer of muscle cells responsible for autonomous phasic contractions, and an outer layer of fibroblasts and a collagen-enriched matrix that connects the vessels to surrounding tissues (Breslin et al., 2019; Bridenbaugh et al., 2013; Petrova and Koh, 2020). In addition, the collecting lymphatic vessels also contain intraluminal valves that prevent backflow and enable unidirectional propulsion of the lymph (Breslin et al., 2019; Zawieja, 2009). Until recently, LMCs were not recognized as a cell type distinct from vascular smooth muscle cells (VSMCs) (Kenney et al., 2020; Muthuchamy et al., 2003). Our data presented here directly show that LMCs are a unique cell type, distinct from other muscle cells including VSMCs.

Dysfunction of the lymphatic system is attributed as the cause of lymphedema and lymphatic diseases, which affect an estimated 7 million patients in the US and 250 million patients worldwide (Rockson, 2018, 2021). Further, there are strong associations for the involvement of the lymphatic system in the pathogenesis of many diseases, including bacterial and filarial infections (Jones et al., 2018; Zhou et al., 2019), inflammatory bowel disease (Rehal et al., 2018), metabolic syndrome (Jiang et al., 2019; Lee et al., 2022; Norden and Kume, 2020), congestive heart failure (Itkin et al., 2021), hypertension (Chachaj et al., 2018), diabetes (Cao et al., 2021) and neurodegenerative diseases (Louveau et al., 2018), including Alzheimer’s disease (Da Mesquita et al., 2018). Further, the recovery from many pathologies, including myocardial infarction (Klaourakis et al., 2021; Liu et al., 2020) and bone regeneration (Biswas et al., 2023), also requires the lymphatic system. Altering the growth and function of the lymphatic system in these diseases and repair processes is a promising therapeutic strategy. Incredibly, there are currently no FDA approved drugs indicated to alter lymphatic function. The discovery of lymphatic vessel-specific therapeutics requires a deep characterization of lymphatic biology that is only now emerging. In the late 1990s and early 2000s, the identification of vascular endothelial growth factor receptor 3 (VEGFR-3) (Kaipainen et al., 1995; Kukk et al., 1996), prospero homeobox 1 (PROX1) transcription factor (Hong et al., 2002; Petrova et al., 2002; Wigle and Oliver, 1999; Wigle et al., 2002), integral membrane glycoprotein podoplanin (PDPN) (Wetterwald et al., 1996), and lymphatic vessel endothelial hyaluronan receptor 1 (LYVE1) (Banerji et al., 1999; Jackson et al., 2001) as critical molecular regulators of LECs revolutionized the field of lymphatic biology research. However, the study of LMCs has fallen behind LECs, in part due to the lack of a comprehensive characterization of LMCs (Kenney et al., 2020). Therefore, it is imperative to improve our knowledge of the molecular signaling in LMCs and their contributions to lymphatic function.

In addition to secondary lymphedemas, many lymphedema patients have congenital primary lymphedemas. Considerable research has not only been dedicated to identifying genes associated with primary lymphedemas to help with diagnosis and to understand the disease pathogenesis, but also to build our understanding of the critical signaling pathways involved in the formation, growth, maturation and function of lymphatic vessels (Brouillard et al., 2021; Mäkinen et al., 2021; Martin-Almedina et al., 2021; Witte et al., 2021). Even with these focused efforts, many cases of primary lymphedema do not have an identified mechanism yet. It is likely that dysfunction of LMC activity and lymphatic contraction are the root cause of lymphedema in some of these patients, as is the case for patients with Cantú syndrome (Davis et al., 2023). The lack of a molecular characterization of LMCs hinders the ability to assign the molecular cause of these primary lymphedemas as well as impairs the ability to discover critical signaling mechanisms necessary for maintaining contractile activity of lymphatic vessel.

Damage to LMCs causes lymphatic pump failure leading to edema and locally immunocompromised tissues (Muthuchamy et al., 2003; Padera et al., 2016; Scallan et al., 2016; Zawieja, 2009). We reported that impaired LMC contraction reduces antigen transportation and is associated with immunosuppression (Liao et al., 2011). Further, we showed that death of LMCs caused by bacterial infection can permanently stop lymphatic pumping (Jones et al., 2018). In addition, functional studies in aged mice and rats have shown decreased LMC coverage of the mesenteric lymphatics (Gashev, 2010), reduced muscle contraction (Akl et al., 2011; Nagai et al., 2011), decreased ion channel protein expression (Zolla et al., 2015), and decreased collecting lymphatic ejection fraction (Akl et al., 2011; Liao et al., 2011; Nagai et al., 2011). Aged mice have decreased lymphatic vessel density, increased lymphatic permeability, and decreased ability to clear bacteria (Kataru et al., 2022). The loss of the contractile phenotype in LMCs during aging causes the reduction of lymph pumping and drainage (Akl et al., 2011), which may help explain the increased infection rate and weakened response to pathogens and vaccines—including COVID-19 vaccines—in older adults (Bartleson et al., 2021).

Single-cell transcriptomics has enhanced our understanding of cell composition and gene expression in lymphatic vessels (Kenney et al., 2022) and lymph nodes (Fujimoto et al., 2020; Rodda et al., 2018; Takeda et al., 2019; Veerman et al., 2019). In this study, leveraging single-cell RNA-sequencing and intravital lymphangiography, we characterized the phenotypes and gene expression plasticity of LMCs in both male and female mice across the lifespan. Our study provides a comprehensive characterization of murine dermal collecting lymphatic vessels and their microenvironment, which will provide the fundamental knowledge to facilitate the study of LMCs in order to identify targets to preserve lymphatic vessel function. Further, identifying the distinct gene signatures of LMCs will provide novel strategies for visualization and cell lineage tracing in collecting lymphatic vessels. Finally, these data will help identify the origins of LMCs during development, the mechanisms driving the regeneration of lymphatic vessels, and the mechanisms involved in various pathophysiological processes, including the onset of primary lymphedema.

## Results

### The single-cell landscape of collecting lymphatic vessels from mouse dermis across lifespan

To characterize the transcriptomes of the cells that make up the walls of collecting lymphatic vessels and the surrounding tissue, we isolated dermal collecting lymphatic vessels that connect the inguinal lymph node and axillary lymph node (**Figure S1**) in 6-week-old (young), 12-month-old (middle-aged), and 18-month-old (aged) C56BL/6J mice from both males and females for 10X Genomics single-cell library construction and subsequent Illumina high throughput sequencing. The total number of reads ranged from 75 million to 100 million in all the samples (**Table 1**). After rigorous quality control (see Methods), 31,140 single-cell transcriptomes passed the quality threshold for downstream analysis. Specifically, we obtained 15,755 single cells in the young group, 5,503 in the middle-aged group, and 9,882 in the aged group (**Table 1**). Of note, the number of cells obtained from male and female mice is comparable, with 14,631 single cells from male mice and 16,509 from female mice (**Table 1**). The average number of genes per cell is between 1000 to 2000 in all samples, and the percentage of mitochondrial genes was less than 5% (**Table 1**), demonstrating the high quality of our single-cell RNAseq datasets (Luecken and Theis, 2019).

Uniform manifold approximation and projection (UMAP) analysis of all the single cells grouped them into 13 cell clusters (Stuart et al., 2019) (**Figure 1A**). Cell type annotations were guided by known marker gene expression (Cohen et al., 2018; Han et al., 2018; Jones et al., 2018; Kalucka et al., 2020; Muhl et al., 2020; Muthuchamy et al., 2003; Petrova and Koh, 2018; Renner et al., 2017; Russell et al., 2022; Siret et al., 2022) (**Figure 1B**) and annotated by age and sex of the mouse of origin (**Figure 1C** and **Table 2**). Our data include a population of *Acta2* and *Notch3* expressing cells that we determined to be LMCs. The presence of large amounts of mesenchymal/fibroblasts, macrophages and epithelial cells in our single-cell data suggests that they are positioned close to the lymphatic vessels and may play a previously unrecognized role in regulating lymphatic vessel function. This is supported by recent data that reported infiltration of tissue-resident macrophages contributes to lymphatic vessel regeneration after injury (Razavi et al., 2023). The CD34+ endothelial cells are likely from adjacent capillaries (Breslin et al., 2019).

**Figure 1.**
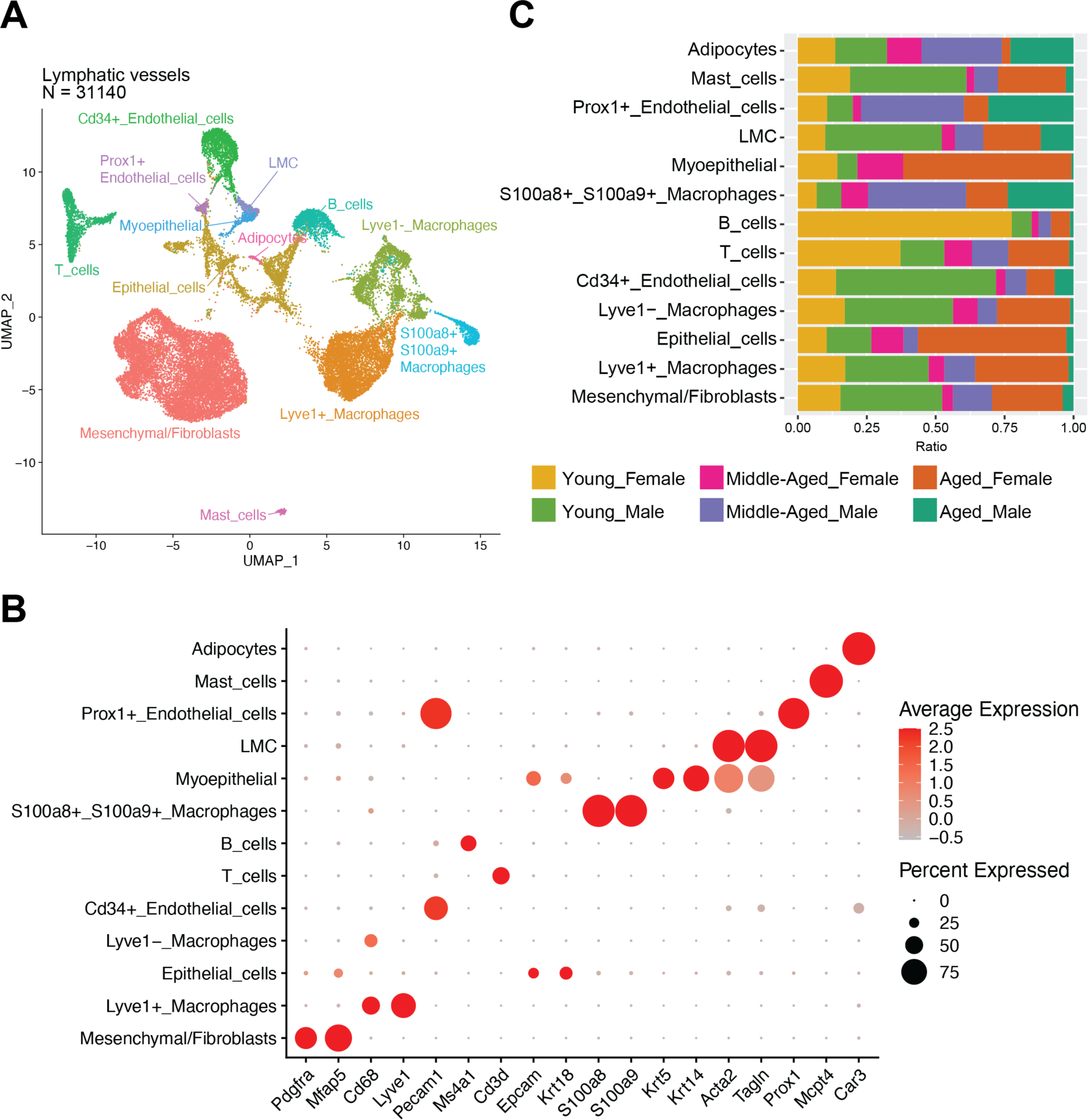
The single-cell atlas of mice collecting lymphatic vessels of various ages. **A**) Uniform manifold approximation and projection (UMAP) embedding of 31,140 cells reveals distinct cell types in the collecting lymphatic vessels and adjacent tissues. The collecting lymphatic vessels between inguinal and axillary lymph nodes were isolated from young (6-week-old), middle-age (12-month-old), and aged (18-month-old) mice of both sexes. The lymphatic vessels from 8-10 mice were combined for single-cell RNA sequencing in each age and sex group. **B**) The dot plot represents the gene expression of cell type-specific marker genes. The size of the circle indicates the proportion of cells expressing the selected genes. Red indicates high expression, while gray indicates low expression. Gene expression values are scaled and log-normalized. **C**) The stacked bar plot represents the proportional distribution of cells across different age and sex groups.

Myoepithelial cells also express the common smooth muscle cell genes *Acta2* and *Tagln*, (**Figure S2A**), but other common marker genes expressed in smooth muscle cells, including *Mustn1*, *Notch3*, *Des*, and *Rgs4* were absent in myoepithelial cells. In addition, *Krt5*, *Krt14*, *Krt17*, and *Cxcl4* were expressed in the myoepithelial cell cluster but not in LMCs (**Figure S2A**). Previous research has reported that myoepithelial cells express *Krt5*, *Krt14*, *Krt17*, and *Cxcl4*, as well as mild levels of *Acta2* and *Tagln* (Han et al., 2018; Kanaya et al., 2019; Karaayvaz et al., 2018), supporting our findings. Of note, 92% of the myoepithelial cells are from female mice (**Figure 1C** and **Table 2**), indicating they are likely to arise from adjacent mammary glands, which are anatomically close to the dermal collecting lymphatic vessels in mice. To confirm this hypothesis, we combined the myoepithelial cells and LMCs from our data with a published single-cell RNA sequencing dataset from murine mammary fat pads (Li et al., 2020). Our analysis showed that LMCs clustered with pericytes, whereas the myoepithelial cells in our dataset clustered with myoepithelial cells in the mammary fat pad dataset (**Figure S2B**).

Overall, we have generated a comprehensive single-cell transcriptomic atlas of murine collecting lymphatic vessels and the adjacent tissues across the lifespan.

### Characterization of the LMC-specific gene signature

Currently, the most commonly employed markers to identify LMCs are αSMA (encoded by *Acta2*) (Jones et al., 2018; Kenney et al., 2020; Russell et al., 2022) and *Myh11* (Muthuchamy et al., 2003). However, these two markers are general markers for smooth muscle cells including VSMCs. Several studies have demonstrated a rich diversity of smooth muscle cells (Liu et al., 2019; Muhl et al., 2022). In addition, it is well-recognized that the LMCs display distinct phenotypes and functions not seen in VSMCs, such as autonomous phasic contractions (Scallan et al., 2016). To identify the unique markers for LMCs, we integrated the LMCs single-cell RNA sequencing data with several seminal studies of muscle cells from different organs, including skeletal muscle cells (GSE143437) (De Micheli et al., 2020), cardiomyocytes (GSE95140, GSE132658, GSE153481) (Goodyer et al., 2022; Nomura et al., 2018; Wang et al., 2020), and vascular smooth muscle cells (GSE203594) (Xu et al., 2022) (**Figure 2A**). We identified a group of genes that are predominantly expressed in LMCs (**Figure 2B**), including *Thbs1*, *Nr4a2*, *Nr4a3*, *Scn3a*, *Adamts4* and *Kcne4*. Of note, *Notch3* has been shown to be a unique marker gene for smooth muscle cells in the heart, lung, and colon, but not in the aorta (Muhl et al., 2022). We also found that LMCs expressed *Notch3* in our single-cell data (**Figure S2A**) and confirmed the presence of Notch3 in LMCs by whole mount immunofluorescence staining (**Figure S3A**). Myosins and actin filaments are the essential components of the muscle cell contractile apparatus (Sweeney and Hammers, 2018). Examination of the single-cell gene expression identified distinct myosins (**Figure 2C-D**) and actin filament molecules (**Figure 2E**) that were expressed in different muscle cells. We found LMCs expressed myosins and actin filaments that are largely similar to aortic smooth muscle cells, including myosin light chain molecules *Mylk*, *Myl9*, *Myl6* (**Figure 2C**), myosin heavy chain molecules *Myh11* and *Myh9* (**Figure 2D**), and actin filament molecules *Actg1, Actb,* and *Acta2* (**Figure 2E**). Cardiomyocytes and skeletal muscle cells showed different expression patterns of actin and myosin molecules (**Figure 2C-E**).

**Figure 2.**
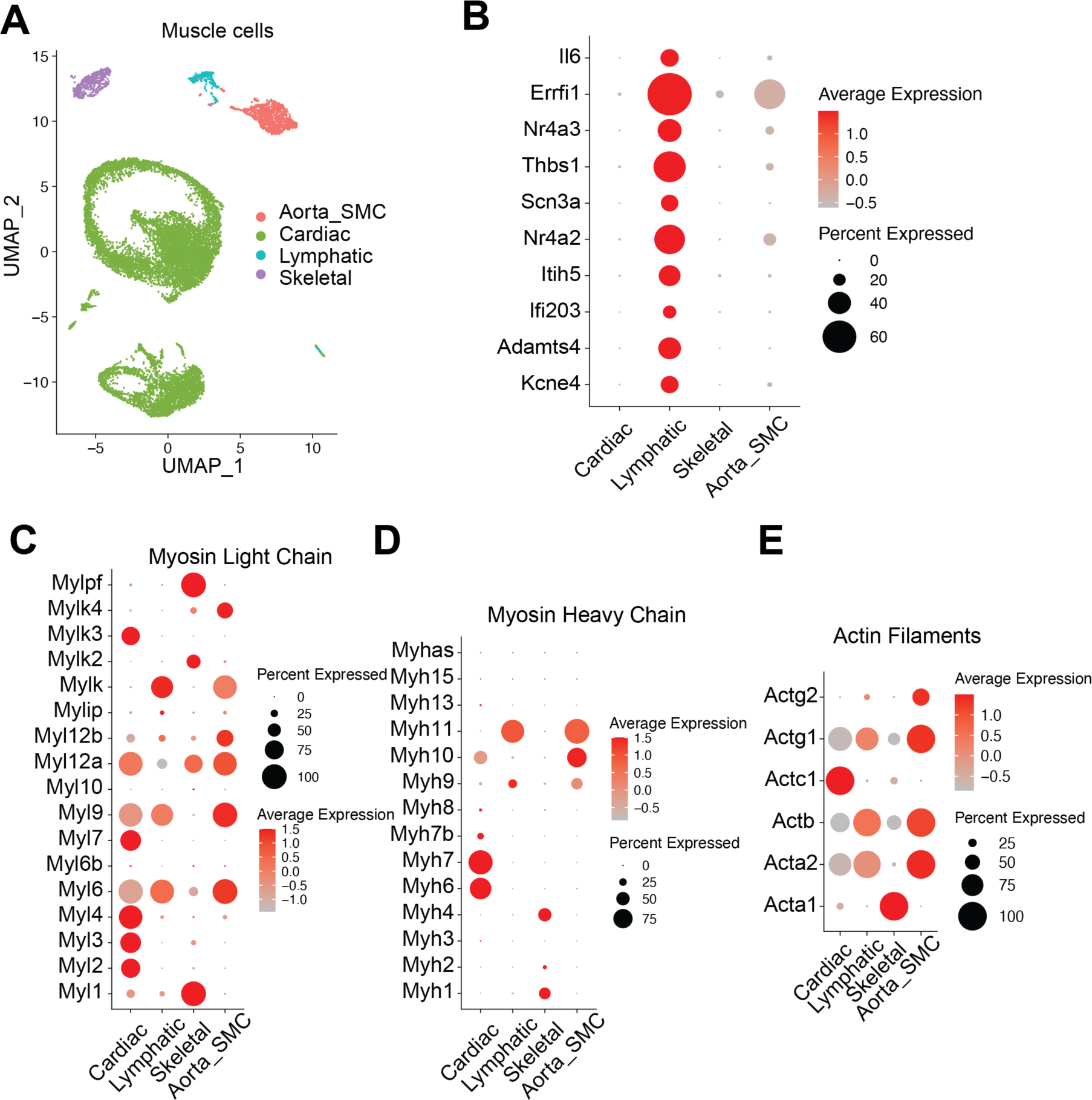
Identify the mouse collecting lymphatic muscle cell-specific marker genes. **A**) UMAP embedding of murine LMCs, aortic smooth muscle cells (GSE203594) (Xu et al., 2022), cardiomyocytes (GSE95140, GSE132658, GSE153481) (Goodyer et al., 2022; Nomura et al., 2018; Wang et al., 2020), and skeletal muscle cells (GSE143437) (De Micheli et al., 2020). **B**) The dot plot displays the single-cell gene expression levels of the top 10 most highly differentially expressed genes in LMCs. **C**) The dot plot displays the gene expression of myosin light chain molecules in all the muscle cells. **D**) The dot plot displays the gene expression of myosin heavy chain molecules in all the muscle cells. **E**) The dot plot shows the gene expression of actin filament molecules in all the muscle cells.

### Differential expression of ion channels in LMCs

Generating electrical action potentials in LMCs creates spontaneous contractions (Hald et al., 2018). Ion channels are necessary to generate action potentials by modulating membrane potential and intracellular calcium concentration (Tykocki et al., 2017). Among these channels, voltage-gated calcium channels (VGCC), voltage-gated potassium (KV) channels, and Ca2+-activated potassium (BK) channels are considered key regulators of vascular smooth muscle excitability and contractility (Pereira da Silva et al., 2022). To examine the presence of ion channels in LMCs, we investigated the expression of calcium, chloride, potassium and sodium channel molecules across different muscle cells.

#### Calcium channels

In contrast to myosin expression where LMCs share similarities with VSMCs, calcium channel expression in LMCs resembled that of cardiomyocytes (**Figure 3A**), including T-type calcium channel genes *Cacna1g* and *Cacna1h* (CaV3.1 and CaV3.2, respectively), and L-type calcium channel genes *Cacna1c* (CaV1.2) (Lee et al., 2014). While L-type calcium channels contribute to the upstroke of the LMC action potential (Telinius et al., 2014a), a recent study showed that T-type calcium channels likely play a minimal or no role in lymphatic pacemaking in mice, despite being present in LMCs (To et al., 2020). Leveraging our intravital lymphatic contraction assay (**Figure S4A**) (Jones et al., 2018; Liao et al., 2011, 2014), we evaluated the effect of calcium channel blockers on lymphatic contraction. We tested the L-type calcium channel blocker (nifedipine) (Lee et al., 2014) and T-type calcium channel blocker (mibefradil) (Lee et al., 2014; To et al., 2020). Of note, at high concentrations, mibefradil can also block L-type calcium channels (Leuranguer et al., 2001). The blockers were administrated either locally [8 μl of nifedipine (17 mg/ml in EtOH) or mibefradil (20 mg/ml)] or systemically [i.p., mibefradil (15 mg/kg)]. The strength of the lymphatic contraction was measured by the ejection fraction—theoretical fractional volume of lymph expelled with each vessel contraction (Jones et al., 2018; Liao et al., 2011, 2014). We found that each of these calcium channel inhibitors can reduce lymphatic contraction after both local and systemic injections (**Figure 3B**). Mibefradil was able to maintain a reduced ejection fraction and contraction frequency for at least 6 hours (**Figure S4B**). These data are consistent with published data in isolated collecting lymphatic vessels (Lee et al., 2014; Telinius et al., 2014a; To et al., 2020), highlighting the importance of calcium channels in lymphatic contractility (Davis et al., 2023b; Russell et al., 2022).

**Figure 3.**
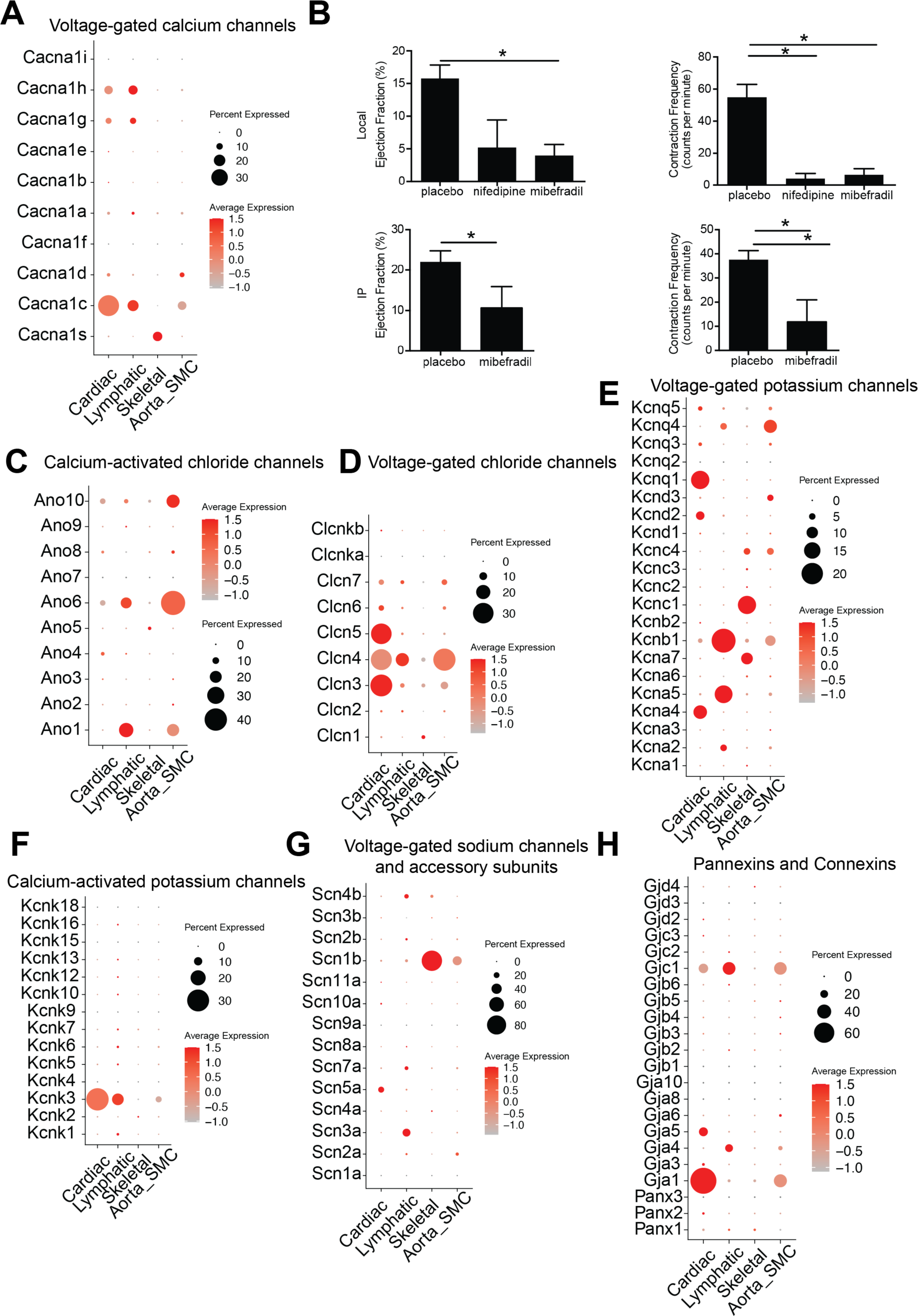
The presence of ion channel molecules in LMCs. **A**) The dot plot shows the gene expression of calcium channel molecules in all the muscle cells. **B**) The lymphatic vessel ejection fraction and contraction frequency after local or systemic administration of calcium blockers. Nifedipine is a L-type calcium channel blocker. Mibefradil is a T-type calcium channel blocker. **C-H**) The dot plot shows the gene expression of C) calcium-activated chloride channels, D) voltage-gated chloride channels, E) voltage-gated potassium channels, F) calcium-activated potassium channels, G) sodium channels and accessory subunits, and H) pannexins and connexins in all the muscle cells.

The release of calcium from the sarcoplasmic reticulum in muscle cells is mediated by ryanodine receptors (RyR) (Felder et al., 2002). Previous studies have reported the presence of RyR in lymphatic vessels and showed that doxorubicin can activate RyR in rat LMCs, causing calcium leak to suppress lymphatic contraction and cause lymphostasis (Stolarz et al., 2019). Dantrolene, a RyR antagonist, prevented doxorubicin-mediated lymphostasis (Van et al., 2021). In line with previous research, we found modest expression of the gene encoding the ryanodine receptor *RyR2* in a small subset of mouse LMCs (**Figure S5A**), consistent with the findings reported in collecting LMCs in rats (Jo et al., 2019).

#### Chloride Channels

The calcium-activated chloride channel, anoctamin-1 (ANO1, gene *Ano1*), also called TMEM16A, contributes to the diastolic depolarization of LMCs and partially sets the lymphatic contraction rate (Zawieja et al., 2019). Consistent with these established findings, we found the presence of *Ano1* mRNA in our murine LMC dataset (**Figure 3C**). Similar to aortic smooth muscle cells, we also find *Ano6* mRNA in LMCs (**Figure 3C**), which has not previously been described. While voltage-gated chloride channels have not previously been described in LMCs, our data show expression of *Clcn4* mRNA in LMCs, as well as in cardiac and aortic muscle cells (**Figure 3D**). Volume-regulated anion channels (VRACs) regulate cell size by passing chloride ions and osmolytes across the cell membrane and have been found to modulate the rate of spontaneous contractions of lymphatic vessels (Solari et al., 2023), but the specific subunits involved are undefined. The genes for the VRAC subunits *Lrrc8a* and *Lrrc8c* are present in our LMC dataset (**Figure S5B**). It is worth noting that the ligand-gated chloride channels were absent in our murine LMC dataset. Similar to cardiac and aortic myocytes, we found LMCs express *Clic1* and *Clic4*—members of the chloride intracellular channel protein family (**Figure S5C**), which has not been previously described in LMCs.

#### Potassium channels

Studies in sheep showed LMC membrane depolarization with exposure to 4-aminopyridine (4-AP), a broadly acting voltage-gated potassium channel inhibitor (Beckett et al., 2007). A previous study showed that the mRNA for the specific *KCNA2* isoform (gene for Kv1.2, a 4-AP sensitive channel) was present in human thoracic duct (Telinius et al., 2014b). In our dataset, we confirm the presence of *Kcna2* mRNA in LMCs and observe some LMCs expressing mRNA for *Kcna5* (protein Kv1.5, critical in atrial myocytes), and *Kcnb1* (protein Kv2.1) (**Figure 3E**). Kv7.4 channels (*Kcnq4*) are not blocked by 4-AP and play a crucial role in the hair cells of the inner ear by participating in potassium recycling, with mutations leading to deafness. Kv7.4 is also found in aortic smooth muscle and in a subset of LMCs in our dataset (**Figure 3E**), which has not been previously described in LMCs.

A *KCNK3* single-nucleotide polymorphism (SNP) was associated with development of secondary lymphedema following breast cancer surgery (Smoot et al., 2017), but neither *Kcnk3* nor any other two-pore domain potassium channels have yet been demonstrated in LMCs (Lee et al., 2022). Our data show *Kcnk3* expression (protein K2P3, also called TASK) in LMCs (**Figure 3F**).

LMCs are known to express *Kcnj8* (ATP sensitive potassium channel Kir6.1) across multiple species and *Kcnj11* (protein Kir6.2) is expressed in some LMC populations (Davis et al., 2022). Cantú syndrome, which has a lymphedema phenotype, can be caused by mutations in the genes *Kcnj8* or *ABCC9* (Davis et al., 2023a). We observed the expression of *Kcnj8*, *Kcnj11*, *Abcc8*, and *Abcc9* genes in a small subset of LMCs (**Figure S5D**). Ether-a-go-go related gene (ERG) potassium channels are key in the repolarization phase of the LMC action potential, and ERG1 (encoded by *KCNH2*) was found to be present in LMCs of multiple species (Kim et al., 2023). We did not find significant mRNA for *Kcnh2* in murine LMCs in our dataset (**Figure S5E**).

#### Sodium channels

Voltage-gated sodium currents were first found in sheep lymphatic muscle (Hollywood et al., 1997). Consistent with a published report in human thoracic duct (Telinius et al., 2015), we found that *Scn3a*, which encodes the sodium channel NaV1.3, is highly expressed in LMCs (**Figure 3G**), Of note, not all the LMCs expressed *Scn3a* (**Figure 3G**), suggesting heterogeneity in the LMC population.

#### Hyperpolarization-activated cyclic nucleotide-gated (HCN) channels

The role of HCN channels in lymphatic pacemaking is controversial (McCloskey et al., 1999; Negrini et al., 2016). Studies have demonstrated that they might play a role in human collecting vessels (Majgaard et al., 2022). We did not find strong mRNA expression of HCN genes in LMCs (**Figure S5F**).

#### Connexins

Connexins have been studied in lymphatic vessels, especially in LECs (Meens et al., 2014). Connexin 37 is required for mouse lymphatic valvulogenesis and development of the thoracic duct (Kanady et al., 2011). In breast cancer patients, mutations in connexin genes correlate with secondary lymphedema following surgery (Hadizadeh et al., 2018). In LMCs, Connexin 45 (gene *Gjc1*) was identified as the critical connexin mediating pacemaker signaling conduction and entrained contractions, with deletions leading to 10-18 fold decrease in conduction speed (Castorena-Gonzalez et al., 2018). We identified the presence of mRNA for *Gja4* (protein: Connexin 37) and *Gjc1* (protein: Connexin 45) in LMCs (**Figure 3H**), confirming existing knowledge about Connexin 45 and suggesting that Connexin 37 may also play a role in LMCs, not just in LECs.

Through the use of freshly isolated collecting lymphatic vessels, our transcriptomic profile of LMCs is consistent with published literature, providing validation of our dataset. Collectively, the LMC transcriptional profile aligns with their phenotypic properties. LMCs can maintain vessel tone similar to VSMCs, yet they can also generate action potential-driven contractions analogous to cardiomyocytes. The distinctive transcriptional activity of LMCs demonstrates similarities to both of these cell types, while distinguishing LMCs as a unique cell type.

### LMC transcription as a function of biological sex

Primary lymphedema is a well-recognized sex-linked disease, with a prevalence that is approximately three times higher in females than in males (Morfoisse et al., 2018, 2021; Smeltzer et al., 1985; Vignes et al., 2021). However, the impact of biological sex on LMC in lymphedema is still poorly understood. We compared the gene expression profiles of LMC between male and female mice and identified 267 differentially expressed genes (**Figure S6A-B**). To gain further insight into the biological consequences of these differentially expressed genes, we performed signaling pathway analysis using PROGENy. Our results showed that estrogen, Trail, MAPK, JAK-STAT, and WNT signaling pathways were up-regulated in the LMC of female mice. In contrast, TGFβ, PI3K, p53, EGFR, TNFα, and NFκB signaling pathways were elevated in the LMC of male mice (**Figure S6C**).

### Lymphatic contraction is impaired and LMC coverage reduced in aged mice

The reduction of mesenteric lymphatic contraction has previously been reported in aged rats (Akl et al., 2011; Nagai et al., 2011) and in peripheral collecting lymphatic vessels in mice (Liao et al., 2011), which is attributed to a reduction of LMC coverage (Gashev, 2010). To examine the impact of aging on lymphatic function, we measured the collecting lymphatic vessel pumping frequency and intensity in young and aged C57BL/6J mice with the intravital fluorescent microscopy (Jones et al., 2018; Liao et al., 2011, 2014). Compared to young mice, the lymphatic vessel contraction strength and frequency in the aged group was significantly decreased (**Figure 4A-B**). Whole mount immunofluorescence staining for αSMA shows that the density of LMCs in aged mice declined substantially (**Figure 4C**). Further, LMCs in the young mice displayed a more circumferential orientation around the vessels compared to LMCs in aged mice, which were more longitudinally oriented along the vessels (**Figure 4C**). The altered orientation of LMCs on the wall in aged lymphatic vessels likely affects their pumping capacity (Kataru et al., 2022).

**Figure 4.**
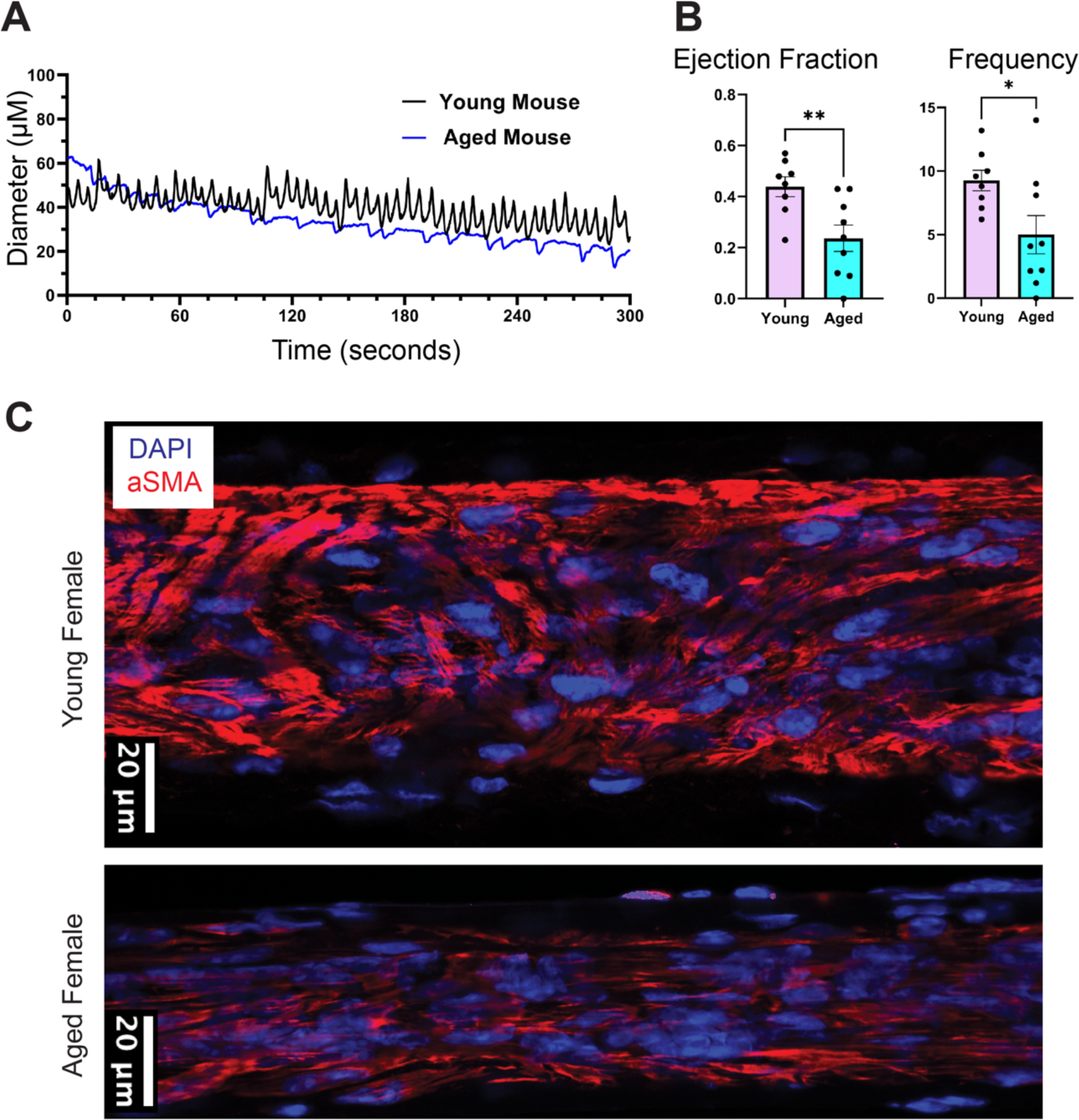
The collecting popliteal lymphatic vessels exhibit diminished contraction and muscle cell coverage during aging. **A)** Representative recording of lymphatic vessel diameter changing over time in a young (6 weeks) and aged (18 months) mouse. **B)** The lymphatic pumping ejection fraction and contraction frequency (cycles/min) in young and aged mice. The ejection fraction for young mice was 0.44 +/- 0.04 (mean +/- SEM), and for aged mice was 0.24 +/- 0.05 (mean +/- SEM), with a mean difference of 0.20 +/- 0.07 (P = 0.008). The frequency for young mice was 9.3 +/- 0.8 (mean +/- SEM, cycles/min), and for aged mice was 5.1 +/- 1.5 (mean +/- SEM, cycles/min), with a mean difference of 4.3 +/- 1.8 cycles/min (P = 0.03). Student’s t-test was used to compare the means between two groups. **C)** Representative whole mount immunofluorescence image of αSMA in lymphatic vessel segments of young (6 weeks) and aged (18 months) C57BL/6 female mice. The scale bar represents 20 μm.

### LMCs transition from a contractile to synthetic phenotype during aging

To understand the transcriptomic alterations in LMCs during aging, we compared the differentially expressed genes between young and aged LMCs. We found 429 differentially expressed genes with a cutoff p-value < 0.05 and fold-change > 0.5 (**Figure 5A**). Gene Set Enrichment Analysis (GSEA) analysis indicates that RNA processing-related gene signatures were up-regulated in the aged LMCs, while supramolecular fiber organization, actin filament-based process, muscle system process, and ion transmembrane transport-related gene signatures were down-regulated in aged LMCs (**Figure 5B**). In addition, we also found that muscle cell contractile apparatus-associated molecules were significantly down-regulated in aged LMCs, including *Acta2*, *Myh11*, *Tpm2*, *Lmod1*, *Col4a1*, *Lamb2*, *Col1a2*, *Col3a1*, *Eln*, and *Col1a1* (**Figure 5C**). The down-regulation of *Acta2* and *Myh11* gene-expression in aged lymphatic vessels was further confirmed by real-time PCR (**Figure 5D**). VSMCs have been shown to be heterogenous, with some displaying a contractile phenotype (enhanced expression of *Acta2*, *Myh11*, *Tagln*, *Myocd*, and *Smtn*) and others displaying a synthetic phenotype (enhanced expression of *Klf4*, *Rbp1*, and *Mmp2*) (Rensen et al., 2007). Our data suggest a similar contractile to synthetic transition occurs in LMCs during aging as shown by the down regulation of contractile associated genes in aged LMCs.

**Figure 5.**
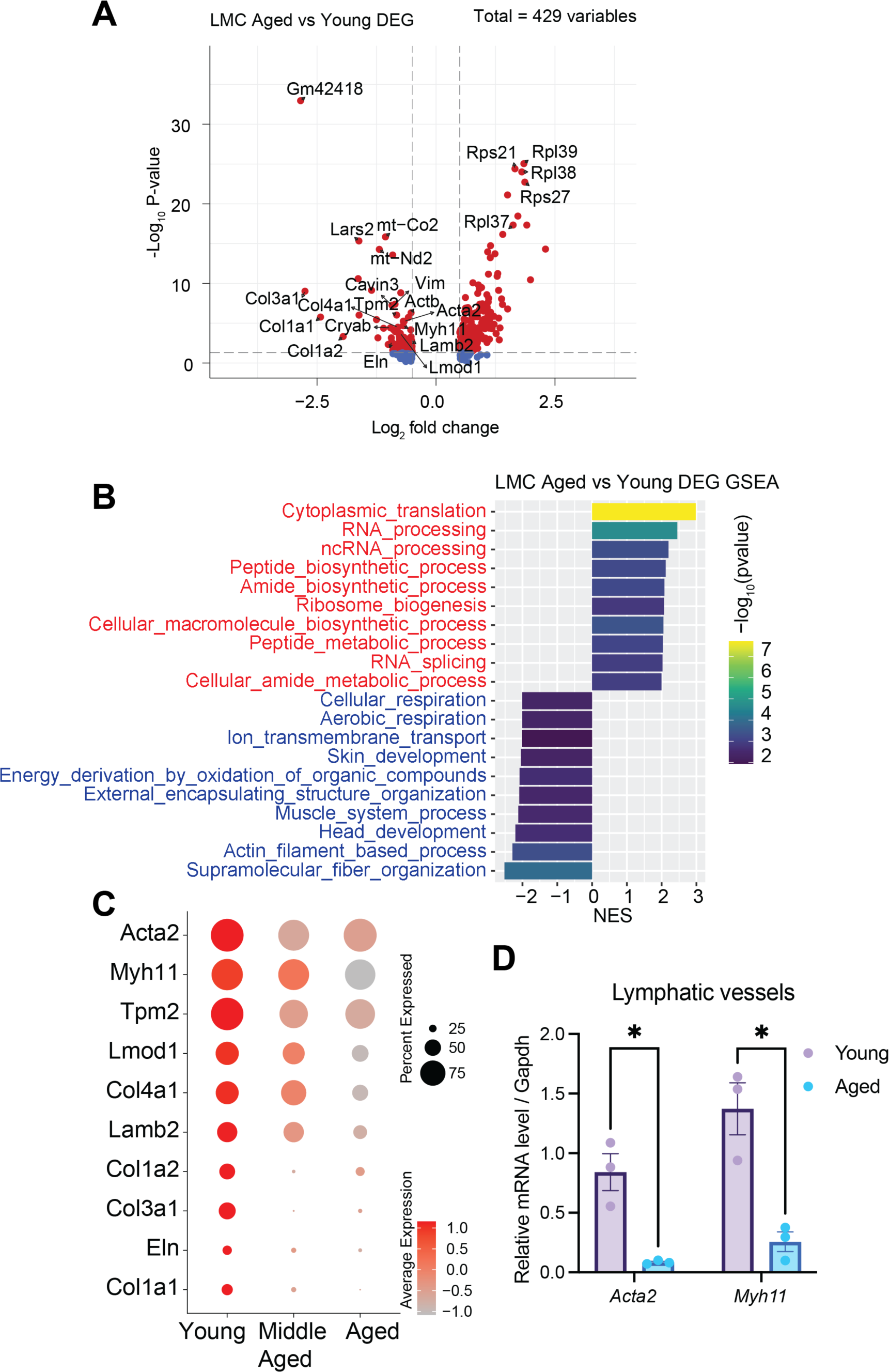
Characterization of murine lymphatic smooth muscle cells during aging. **A)** The volcano plot represents the differentially expressed genes in LMCs between the young (6-week-old) and aged (18-month-old) groups. The red dots indicate significant differentially expressed genes with a fold change > 0.5 and P value <0.05. **B)** Gene set enrichment analysis (GSEA) of significant differentially expressed genes in LMCs between young (6-week-old) and aged (18-month-old) mice. The gene ontology biological process gene sets were used as a reference for the GSEA analysis. The gene sets are ordered by normalized enrichment score (NES), and the bar plot displays the top 10 up- or down-regulated gene sets. Positive values of NES show higher expression in aged LMCs, and negative values of NES show higher expression in young LMCs. **C)** The dot plot represents the gene expression of muscle cell contractile-associated genes in LMCs. The size of the circle indicates the proportion of cells expressing the selected genes. Red indicates high expression, while gray indicates low expression. Gene expression values are scaled and log-normalized. **D**) The gene expression of *Acta2* and *Myh11* in collecting lymphatic vessels in young (6-week-old) and aged mice (18-month-old) was quantified by real-time PCR. In each group, the lymphatic vessels were isolated from one male mouse and one female mouse, which were combined for RNA extraction. Student’s t-test was used to compare the means between two groups. *p<0.05.

### TNFα can drive the contractile to synthetic switch of LMCs in aged lymphatic vessels

Previous studies suggest that the cells in lymphatic vessel adjacent tissue impact the contraction of LMCs by releasing signaling molecules, such as nitric oxide (Chen et al., 2017; Liao et al., 2011; Rehal et al., 2020). To understand whether the transition of the LMC phenotype from contractile to synthetic during aging was attributed to paracrine signals from the adjacent tissue, we analyzed cell-cell interactions of each signaling pathway in the single cell RNA sequencing datasets from young and aged lymphatic vessels to identify the conserved and context-dependent signaling pathways. We found that MK, NCAM, GRN, CD200, CD23, IL4, and MPZ cell-cell communication probabilities were greater in the lymphatic vessels in young mice (**Figure S7A)**. In contrast, the TGFβ, ANNEXIN, ITGAL-ITGB2, FGF, IL6, BMP, EPHB, NECTIN, WNT, NRXN, EPHA, GDF, VISFATIN, and TNF cell-cell communication probabilities were significantly enriched in the aged lymphatic vessels (**Figure S7A**). To examine whether these signals were directly targeting LMCs, we performed ligand-receptor communication probability analysis (Jin et al., 2021). We found that Tnf-Tnfrsf1a, Igf-Igf1r, Fgf2-Fgfr1, and Bmp2-(Bmpr1a+Bmpr2) ligand-receptor interactions were significantly up-regulated in aged LMCs (**Figure 6A**). The cell-cell interaction network analysis of the TNF signaling pathway indicated Lyve1+ macrophages, S1008a+ S100a9+ macrophages and mast cells were the major cell types that frequently interact with LMCs in aged mice (**Figure 6B**). Single-cell gene expression analysis also showed a high level of *Tnf* mRNA in Lyve1+ macrophages, S1008a+ S100a9+ macrophages and mast cells in aged mice (**Figure 6C**). To test if TNFα can induce a contractile to synthetic transition in LMCs, we isolated LMCs from young mice and cultured the LMCs *in vitro* in the presence or absence of TNFα (10 ng/mL) (Arroyo-Ataz and Jones, 2022). We found that TNFα suppressed the gene expression of *Myh11* and *Acta2* (**Figure 6D**). Taken together, we hypothesize that in aged lymphatic vessels, TNF-α derived from macrophages and mast cells suppresses the contractile apparatus in LMCs, resulting in a reduction of lymphatic contraction (**Figure 6E**).

**Figure 6.**
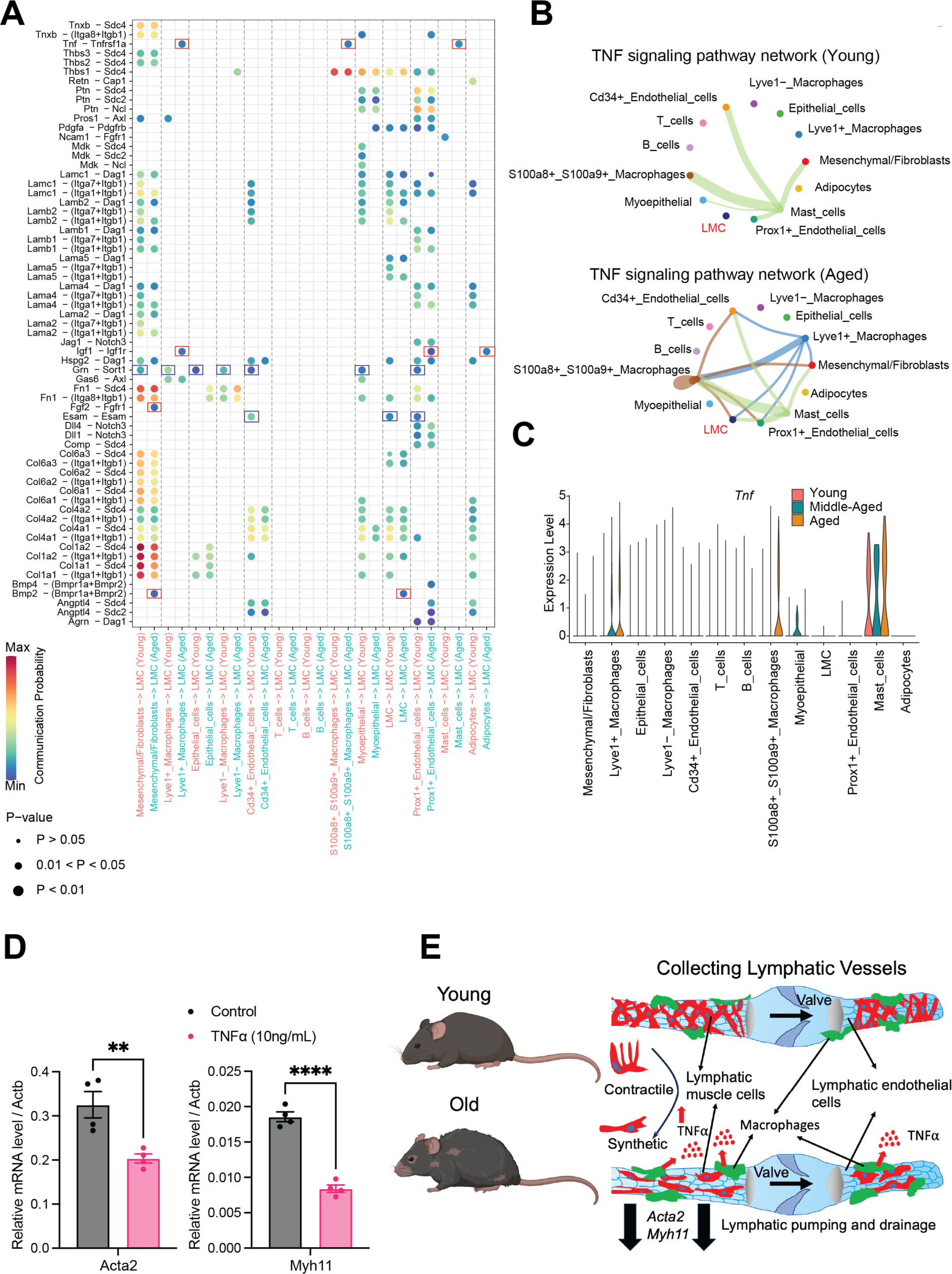
The cell-cell interactions between cells that comprise lymphatic vessels and cells in adjacent tissues. **A**) Dot plot describing the ligand-receptor interactions between cells of the lymphatic vessels and adjacent tissue (source) and muscle cells (target) in young (6-week-old) and aged (18-month-old) mice. The size of the circle represents the p-value of the communications. The color of the dot represents the communication probability of the ligand-receptor pair between cells. Blue represents low probability, and red represents high probability. The red box highlights the ligand-receptor pairs that were elevated in the aged (18-month-old) mice, blue box highlights the ligand-receptor pairs that were decreased in the aged (18-month-old) mice. **B**). The network of cell-cell communication in TNFα signaling pathway in young (6-week-old) and aged (18-month-old) mice. **C**) The violin plot shows the gene expression of *Tnf* in all the cells of various ages. **D**) The gene expression of *Acta2* and *Myh11* in lymphatic muscle cells after TNFα treatment. The lymphatic muscle cells were collected from the afferent lymphatic vessel to the popliteal lymph node of young (6-week-old) mice and cultured in the presence of 10 ng/mL TNFα for 48 hours. **E**) Macrophages and mast cells positioned in the microenvironment of aged lymphatic vessels produce TNFα, which suppresses the gene expression of *Acta2*, *Myh11* and other contractile apparatus-related genes and drives the contractile to synthetic transition of LMCs. The reduction of LMC coverage and decreased contractile phenotype in aged lymphatic vessels lead to a lower ejection fraction and contraction frequency. ** p< 0.01; **** p<0.0001.

## Discussion

With hundreds of millions of patients around the globe suffering from lymphedema and lymphatic diseases (Rockson, 2018, 2021), our therapeutic tools to help these patients are severely lacking, making these patients some of the most underserved by biomedical research. The lymphatic system covers the surface of the brain (Louveau et al., 2015) and is in the tips of the toes and is in every organ system in between, yet there are no FDA approved drugs indicated to alter its function. One limitation is the lack of fundamental knowledge of molecular phenotypes of all the cell types necessary to build and drive the function of lymphatic vessels. Rapid progress has been made in characterizing the critical lymphatic endothelial cell (Alitalo, 2011; Aspelund et al., 2016; Baluk et al., 2007; Ducoli and Detmar, 2021; Hong et al., 2002; Oliver et al., 2020; Takeda et al., 2019; Wigle et al., 2002), but limited recognition of LMCs as a unique cell type, distinct from VSMCs, has hindered progress. Due to their important functional roles in driving lymph flow, LMCs are important targets for potential therapeutics. Here, we provide a comprehensive characterization of the LMC transcriptomic output across the lifespan, accounting for biological sex (**Figure 1A&C**). Our data directly show that LMCs are a unique muscle cell type with similarities in transcriptional activity to some aspects of both VSMCs and cardiomyocytes (**Figure 2C-E and Figure 3A**), consistent with the functional activity of LMCs in driving lymphatic pumping (Scallan et al., 2016; Zawieja, 2009). This dataset provides a resource that can spur discovery of targeted therapeutics to alter lymphatic vessel contractility by altering activity of LMCs or restoring lost LMC activity that occurs with aging. Further, this data resource can be used to help link genetic mutations associated with human primary lymphedemas to testable functional consequences for the lymphatic system. Through newly uncovered connections in primary lymphedema, further discovery of fundamental lymphatic biology can occur as well as the potential to develop therapies for patients suffering from these primary lymphedemas.

Through single-cell RNA sequencing analysis, we uncovered a list of LMC marker genes (**Figure 2B**), including *Thbs1*, *Nr4a2*, *Nr4a3*, *Scn3a*, *Adamts4* and *Kcne4*. In addition, we discovered that LMCs express myosin heavy and light chain as well as actin filament molecules similarly to VSMC (**Figure 2C-E**). This is supported by a recent study that utilized Cre-tdTomato reporter mice specific for progenitors of skeletal muscle cells (Pax7^+^ and MyoD^+^) and VSMCs (Prrx1^+^ and NG2^+^) to investigate the origin of LMCs in lymphatic vessels afferent to the popliteal lymph node (PLV-LMCs) (Kenney et al., 2020, 2023). The study demonstrated that PLV-LMCs originate from Pax7^−^/MyoD^−^/Prrx1^+^/NG2^+^ progenitors—similarly to VSMCs—prior to postnatal day 10. However, after postnatal day 10, PLV-LMCs originate from a previously unknown Pax7^−^/MyoD^−^/Prrx1^+^/NG2^−^ muscle progenitor.

Voltage-gated calcium channels transduce membrane depolarization into augmented Ca2+ influx into cells and play an important role in the regulation of both contraction and gene expression in vascular smooth muscle cells (Pereira da Silva et al., 2022; Tykocki et al., 2017), cardiomyocytes (Marbán, 2002), and LMCs (Lee et al., 2014; To et al., 2020). The intracellular calcium channel serves as a phase controller by regulating the myosin-light-chain kinase activity (Kim et al., 2008). Depolarization of LMCs activates and opens L-type voltage-gated calcium channels and mediates entry of Ca2+ into LMCs, which is crucial for excitation-contraction coupling (Lee et al., 2014; To et al., 2020). As expected, we found both L-type and T-type voltage gated calcium channels in LMCs (**Figure 3A**). In addition, our functional data confirms that the calcium channel blockers nifedipine and mibefradil decrease lymphatic contraction frequency and strength (**Figure 3B**). Ryanodine receptors in the sarcoplasmic reticulum also contribute to calcium release and electromechanical coupling (Stolarz et al., 2019; Van et al., 2021). Our findings indicate that LMCs express RyR2 (**Figure S5A**), suggesting a potential unrecognized role in regulating lymphatic contraction.

The voltage-gated sodium channel, Nav1.5, is crucial for generation of the cardiac action potential, and vascular smooth muscles lack voltage-gated sodium channels. Previous studies indicate that the contribution of voltage-gated sodium channels to the lymphatic action potential may vary by species (Chan and von der Weid, 2003; Ohhashi et al., 1980; Telinius et al., 2015). Tetrodotoxin, a blocker of several types of voltage-gated sodium channel sensitive currents are found in human (Telinius et al., 2015) and sheep, but not in bovine (Ohhashi et al., 1980) or guinea-pig (Chan and von der Weid, 2003) lymphatic vessels. We find that Nav1.3 mRNA is present in murine LMCs (**Figure 3G**).

In addition to voltage-gated sodium channels, multiple other channel types, such as Ano1, HCN, and other cationic channels may contribute to the pacemaking and upstroke of lymphatic depolarization. Different from rat (Negrini et al., 2016), sheep (McCloskey et al., 1999) and human (Majgaard et al., 2022) lymphatic vessels—which have HCN-mediated currents—the mRNA expression for HCN was minimal in our murine LMC dataset (**Figure S5F**). *Ano1* was present in our dataset (**Figure 3C**), consistent with its known role in LMC depolarization (Zawieja et al., 2019). Additionally, the transcript for another calcium-activated chloride channel, *Ano6*, was also present (**Figure 3C**), although it has not been previously studied in LMCs. We also identified that LMCs expressed chloride channel genes *Clic1* and *Clic4* (**Figure S5C**), which have not been studied in LMCs. After depolarization and initiation of LMC twitch contraction, the cells must repolarize to reset the resting membrane potential. In cardiomyocytes, repolarization is driven by a multitude of potassium channels (Grant, 2009), and the same has been presumed for LMCs (Kim et al., 2023). Consistent with existing LMC literature, we found the following potassium channel genes: *Kcna2* (Kv1.2), *Kcnj8* (Kir6.1), *Kcnj11* (Kir6.2), *Abcc9* (SUR2), *Kcnh2* (ERG1) (**Figure 3E, S5D-E**).

Additional transcripts for potassium channels identified by our dataset which have not previously been studied in LMCs includes: *Kcna5* (protein Kv1.5), *Kcnb1* (Kv2.1), *Kcnq4* (Kv7.4), *Kcnk3* (K2P3), and *Abcc8* (SUR1) (**Figure 3E-F, S5D**). In addition, the voltage-gated potassium (Kv) channel β subunit gene *Kcne4*, is expressed in LMCs, but not in cardiomyocytes, skeletal muscle or VSMCs (**Figure 2B)**. KCNE4 is known to inhibit the current of Kv2.1 and Kv7.4 (David et al., 2015), both of which are expressed on LMCs (**Figure 3E**). Electrically driven contraction mediated by LMCs needs to be propagated along a lymphangion for coordinated systole in a lymphatic vessel. The gap junction protein connexin 45 is involved in primary lymphedema, has been found to mediate entrainment of LMC contraction waves (Castorena-Gonzalez et al., 2018), and is also present in our dataset (*Gjc1*) (**Figure 3H**). In addition, we also identified the presence of *Gja4* (connexin 37) in LMCs (**Figure 3H**).

Aging is a natural process that affects all tissues and organs, including lymphatic vessels. As the body ages, oxidative stress can accumulate in lymphatic vessels (Thangaswamy et al., 2012). Mechanistically, the accumulation of reactive oxygen species and pro-inflammatory cytokines— such as tumor necrosis factor-alpha (TNFα), interleukin-1beta (IL-1β), interleukin-6 (IL6)—can promote the activation of “sheddases”, including metalloproteinases, heparinase, and hyaluronidase (González-Loyola and Petrova, 2021). This pro-inflammatory microenvironment triggers the accumulation of sheddases and accelerates glycocalyx degradation, leading to vascular hyperpermeability, unregulated vasodilation, microvessel thrombosis, and increased leukocyte adhesion (González-Loyola and Petrova, 2021). Previous studies demonstrated that aging increases the proliferation and migration of VSMCs and accelerates the contractile to synthetic transition of these cells (Bochaton-Piallat et al., 2001; Rensen et al., 2007). Similarly, we also found the reduction of gene-expression of contractile-related genes in aged LMCs, which can be driven by the inflammatory cytokine TNFα that increases in tissues with age. Consequently, the phenotypic change of LMCs led to the attenuation of lymphatic pumping and drainage (**Figure 4A-B**). Further, the loss of muscle cells in lymphatic vessels can lead to a decrease in contraction frequency, systolic lymph flow velocity, and pumping activity, ultimately impairing lymph flow (Jones et al., 2018; Zolla et al., 2015). Consequently, age-related lymphatic impairment can cause various disorders, including imbalances in interstitial fluid homeostasis, abnormal lipid accumulation, and impaired immune responses to infections, cancers, and vaccines.

LMC-disruption has been linked to pathological conditions such as lymphedema (Scallan et al., 2016), which is characterized by swelling due to inadequate tissue interstitial fluid absorption and lymph transport. Lymphedema profoundly impacts a patient’s life by limiting limb or organ function and making the patient prone to infections that often require hospitalization. Unfortunately, current treatments are cumbersome and often inadequate (Rockson, 2018, 2021). There is no cure for lymphedema. Our preclinical animal models have demonstrated that methicillin-resistant *Staphylococcus aureus* (MRSA), bacteria that often causes skin and soft tissue infections, can damage LMCs and stop lymphatic pumping and lymph flow (Jones et al., 2018). Persistent MRSA infections can progress to sepsis and individuals aged 65 years and older are particularly susceptible to sepsis (Lee et al., 2018; Moreira et al., 2021). Of note, the deficits in LMC coverage of lymphatic vessels persists even after the infection resolves, leading to questions of why LMC do not functionally regenerate (Jones et al., 2018). Determining the origin of LMCs and their developmental signaling will provide new strategies to regenerate LMCs on damaged lymphatic vessels. While the origins of LECs have been widely studied (Butler et al., 2009; Grimm et al., 2023; Gutierrez-Miranda and Yaniv, 2020; Hong et al., 2002; Petrova et al., 2002; Wigle and Oliver, 1999; Wigle et al., 2002), the origin and development of LMC remain largely unknown. In addition, whether or not LMC stem cells exist in adults remains an open question in the field.

A recent study has identified markers of cellular heterogeneity in subsets of smooth muscle cells across multiple organs (Muhl et al., 2022). In another study, Kenney et al. characterized the muscle cells in popliteal lymphatic vessels in wild-type, Cspg4-Cre/Ai9-tdTomato, and TNF-Tg mice using fluorescence-activated cell sorting (FACS) for single-cell RNA sequencing (Kenney et al., 2022). In both studies, the cells were sorted with flow cytometry before single-cell RNA sequencing, which is known to alter the transcriptome of cells. In contrast, we generated droplet-based single-cell RNA sequencing profiles from all the cells in freshly isolated lymphatic vessels and adjacent tissues without presorting (**Figure 1A**). Our unbiased approach also allows the analysis of cell types previously lost to cell sorting. As we have shown in our data, *Acta2* and *Tagln* are also expressed in myoepithelial cells (**Figure 1B, S2A**), suggesting that the previous cell sorting-based approach could introduce bias to the datasets if they relied on αSMA to identify cells. Characterization of cell types with a few marker genes does not capture the spectrum of functional states and gene expression programs in lymphatic vessels. Our data show that by surveying all cells in the mouse dermal collecting lymphatic vessels, we identified markers of cellular heterogeneity and distinct cellular subpopulations that show plasticity with aging. In conclusion, we were able to utilize single-cell RNA sequencing to uncover the identity of LMCs and to analyze differential gene expression in LMCs across the murine lifetime (**Figure 5A-C**). Our findings provide markers of LMCs which will aid in their future study and may help associate gene mutations identified in patients with primary lymphedema with a functional deficit in their lymphatic system. In addition, we identified TNFα as a potential mechanism (**Figure 6A-D**) to help understand changes in lymphatic vessel function during aging.

### The limitations of this study

In this study, we focused on the collecting lymphatic vessels in dermis, which often causes lymphedema in humans when damaged. It is possible that LMCs in mesenteric lymphatics and the thoracic duct may exhibit different gene signatures (Zawieja et al., 2018). Emerging data also suggest that lymphatic vessels exhibit organ-specific heterogeneity (Oliver et al., 2020; Petrova and Koh, 2018) and acquire distinct characteristics upon exposure to different tissue microenvironments (Aspelund et al., 2016; Biswas et al., 2023; Da Mesquita et al., 2018; Reed et al., 2019), which in turn enable them to execute multifaceted functions. Characterizing organ-specific lymphatic vessels and LMCs will help gain insight into their heterogeneity and specialization, enabling the specific targeting of organ-associated vessels for therapeutic purposes. In addition, our single-cell data is from mice and human LMCs may display different gene signatures.

### Methods and materials

#### Mice

C57BL/6 mice were housed in animal facilities at Massachusetts General Hospital. Both male and female mice of ages 6 weeks, 12 months, and 18 months were utilized for this study. The 18-month-old mice were provided by the National Institute on Aging/National Institutes of Health. All animal procedures were subjected to review and approval by the Institutional Animal Care and Use Committees (IACUC) at Massachusetts General Hospital.

#### Collecting lymphatic vessel isolation

Mice were anesthetized with ketamine (100 mg/kg) and xylazine (10 mg/kg) via intraperitoneal injection. The ventral fur was removed via shaving and the surgical site was disinfected with 70% ethanol. A ventral midline skin incision was made from the pubis to the sternum. Lateral incisions were extended from the midline to the upper and lower limbs. The skin was then opened laterally to expose the collecting lymphatic vessels that connect the inguinal lymph node and axillary lymph node (inguino-axillary lymphatic vessels). To enhance the visibility of the lymphatic vessels, 4 μL of Evans Blue (0.1% diluted in PBS) was injected into the inguinal lymph node and filled the lymphatic vessels from the inguinal to the axillary node. The collecting lymphatic vessels were isolated under a stereo microscope, and excess fat and connective tissues were removed while the vessels remained in situ within the mouse to maintain the health of the tissue during the surgical preparation of multiple mice. Once the vessels were cleaned from all mice, they were cut and removed from each mouse. The isolated lymphatic vessels were pooled and immediately transferred to a Petri dish containing high-glucose DMEM (Corning 10-013-CV) without FBS and kept on ice until digestion. Throughout the dissection, mice were kept under anesthesia to minimize pain and discomfort.

#### Single-cell RNA-Seq library preparation

Eight to ten mice were used in each of six groups (two sexes: male, female; and three ages: 6 weeks, 12 months, 18 months). The harvested lymphatic vessels were transferred into 1.5 mL Eppendorf tubes containing 1 mL of digestion buffer [10 mL high glucose Dulbecco’s Modified Eagle Medium (DMEM) (Corning 10-013-CV) with 20 mg Collagenase I, 5 mg Elastase, 8 mg Dispase, and 1 mg DNaseI]. The vessels were incubated in a 37°C swirling water bath for 40-60 minutes, with the digestion contents being gently resuspended every 5 minutes. After digestion, the samples were filtered through a 70 μM nylon mesh and quenched by adding fetal bovine serum (FBS) to a final concentration of 10%. The samples were then centrifuged at 300 rcf for 5 minutes at 4°C. The supernatant was discarded, and the pelleted cells were resuspended in 1 mL of ACK lysis buffer and kept at room temperature for 2 minutes to remove the red blood cells. The ACK lysis reaction was quenched by adding 9 mL of FACS buffer (2% FBS + 1X DPBS). The resulting 10 mL sample was centrifuged at 300 rcf for 5 minutes at 4°C. The pelleted cells were resuspended in 150-200 μL of FACS buffer and filtered through a 35 μM nylon cell strainer. Cell viability was confirmed by Trypan Blue (Gibco) staining through manual counting as well as TC20 automated cell counting (Bio-Rad). Only samples with > 74% live cells were processed with 10X Genomics Single Cell 3’ Reagent Kit-based Gel Bead-In-Emulsions (GEM) reactions, followed by Illumina NextSeq 2000 sequencing with the High Output PE26/98 mode.

#### Single-cell RNA-Seq data analysis

The Cell Ranger analysis pipeline was utilized to pre-process and align raw fastq-format data and generate feature-barcode matrices (Zheng et al., 2017). Subsequently, Seurat was employed for graph-based single-cell clustering (Stuart et al., 2019). To ensure the selection of high-quality single-cell transcriptomic profiles, cells that had less than 100 genes or more than 10,000 featured RNAs detected or more than 25% mitochondrial genes, were excluded. Genes expressed in less than 3 cells were also removed. The R package SCpubr was used for single-cell RNA sequencing data visualization, pathway analysis, and transcription factor analysis (Blanco-Carmona, 2022). Cell cycle regression was performed based on the Seurat cell cycle scoring and regression method. Gene ontology functional annotation was performed using the ShinyGO software (Ge et al., 2020). Additionally, the CellChat R package (Jin et al., 2021) was used for cell-cell interaction network and ligand-receptor analysis, with the CellChatDB mouse database manually curated based on literature-supported ligand-receptor interactions. The R packages clusterProfiler (Wu et al., 2021) and msigdbr (Liberzon et al., 2015) were employed for Gene Set Enrichment Analysis.

#### RNA extraction and real-time PCR assay from lymphatic vessels

The collecting lymphatic vessels in 6-week-old and 18-month-old mice were isolated as described above, and immediately placed in 1.5 mL Cryogenic vials (ThermoFisher Scientific) and kept in liquid nitrogen to preserve the RNA quality. PicoPure RNA Isolation Kit (ThermoFisher Scientific, Cat. #: KIT0202, KIT0204) was used for RNA extraction based on the manufacturer’s instructions. After RNA extraction, iScript cDNA Synthesis Kit (BioRad) was employed for the cDNA synthesis. Two mice (4 inguino-axillary lymphatic vessels) were used per RNA extraction reaction. The cDNA from multiple extraction reactions was pooled in a single cDNA master tube. Next, real-time PCR for *Myh11* and *Acta2* was performed according to the manufacturer’s instructions using iTaq Universal SYBR Green Supermix (Bio-Rad, Cat. #: 1725124) on a Bio-Rad CFX Connect Real-Time PCR Detection System.

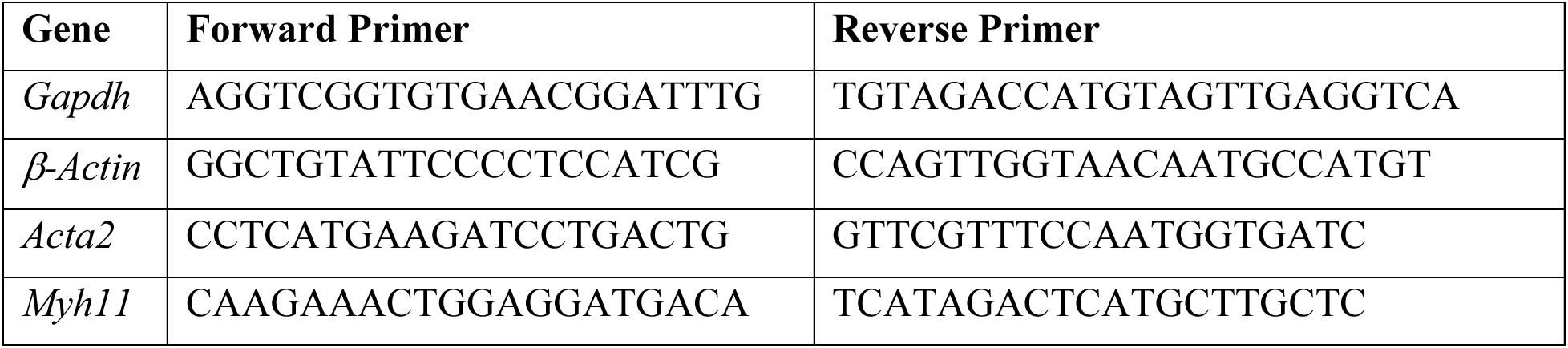

#### Staining of whole mount collecting lymphatic vessels

For αSMA whole mount staining, inguino-axillary lymphatic vessels or the afferent collecting lymphatic vessels to the popliteal lymph node were collected as described above. Next, the isolated vessels were fixed for two hours at room temperature in 4% formaldehyde, washed three times in 1x PBS for 15 minutes, then stored in 1x PBS at 4°C for overnight. The following day, the lymphatic vessels were blocked and permeabilized for 3 hours at room temperature in 10% normal donkey serum (NDS) and 0.1% Triton-X in PBS. After that, lymphatic vessels were washed in 1x PBS three times, with each wash lasting 5 minutes. Next, lymphatic vessels were carefully cut into smaller pieces and stained with αSMA-Cy3 (1/200 dilution, Clone: 1A4, MilliporeSigma, Cat. #: C6198) and DAPI in 1x PBS with 1% NDS and 0.05% Triton-X at 4°C for overnight. The following day, lymphatic vessels were washed in 1x PBS at room temperature three times, with each wash lasting 15 minutes. After that, the lymphatic vessels were mounted onto glass slides using ProlongGold liquid mountant and glass coverslips. Slides were cured flat on the benchtop at room temperature for at least 48 hours prior to imaging. Samples were imaged using a Zeiss LSM 880 Airyscan confocal microscope with 40x oil objective at one Airyunit.

For Notch3 and αSMA wholemount staining, the afferent collecting lymphatic vessels to the popliteal lymph node were collected from Prox1-GFP mice as described above. The popliteal lymphatic vessels were fixed with 4% PFA for 30 minutes, washed with 1x PBS, and incubated with antibodies and DAPI overnight at 4°C (Notch3-APC, 1/50 dilution, Clone: HMN3-133, Biolegend, Cat:130512; aSMA-Cy3, 1/200 dilution, Clone: 1A4, MilliporeSigma Cat: C6198). The following day, lymphatic vessels were washed three times in 1x PBS, with each wash lasting 5 minutes. The tissue was then placed on a glass slide to visualize lymphatic vessels using a confocal microscope (Olympus 1X81) with 4x, 10x, and 20x air objectives, and 60x oil objectives (Olympus). The raw image data in multi-tiff format were processed by ImageJ.

#### Intravital lymphatic pumping assay

Mice were anesthetized using ketamine (100 mg/kg) and xylazine (10 mg/kg). 3 μL of 2% FITC-Dextran (2 million molecular weight) in 0.9% normal saline was injected intradermally into the footpad. The surgical site was prepared by carefully removing the skin and underlying connective tissue near the afferent lymphatic vessel leading to the popliteal lymph node (PLV) as described previously (Jones et al., 2018; Liao et al., 2011, 2014). Time-lapse imaging was performed using an inverted fluorescence microscope (Olympus Ix70) to record 2250 images at 133 ms intervals. Custom MATLAB code was employed for image analysis. The vessel wall position was tracked, and frequency and ejection fraction was measured using a peak-and-valley method as described previously (Liao et al., 2014). Ejection fraction serves as a measure of the strength of lymphatic contraction.

#### In vitro assay of lymphatic muscle cell response to TNFα

The collecting lymphatic vessels in young mice (6-week-old) were isolated according to previously described methods with slight modifications (Arroyo-Ataz and Jones, 2022). Briefly, mice were anesthetized with ketamine (100 mg/kg) and xylazine (10 mg/kg) followed by a footpad injection of 4 μL 4% Evans Blue Dye to visualize the lymphatic vessels. After that, the surgical site was shaved and prepared by removing the skin and underlying connective tissue to expose the afferent lymphatic vessel to the popliteal lymph node. The isolated lymphatic vessels were digested for 30 minutes at 37°C with agitation in an enzymatic cocktail consisting of 1 mg/mL collagenase I (Worthington, Cat. #: LS004196), 1 mg/mL Dispase II (Gibco, Cat. #: 17105041), 100 μg/ml DNAse I (Millipore Sigma, Cat. #: 04716728001) and 200 μg/ml Liberase (Millipore Sigma, Cat. #: 05401020001). The digestion was quenched with 10% FBS and resuspended in 20% FBS DMEM (0.5 ml per animal). Cells were plated in a 24-well plate and grown for 4-5 days, after which they were treated with TNFα (Biolegend, Cat. #: 575202) for 24-48 hours. Cells were pelleted and then resuspended in RLT lysis buffer. RNA was extracted using a Qiagen RNeasy kit (Qiagen, Cat. #: 74004) based on the manufacturer’s instructions. Verso cDNA Synthesis Kit (Thermo Scientific, AB1453A) was employed for cDNA synthesis. Next, real-time PCR for *Myh11* and *Acta2* was performed according to the manufacturer’s instructions using iTaq Universal SYBR Green Supermix (Bio-Rad, Cat. #: 1725124) on a Bio-Rad CFX Connect Real-Time PCR Detection System.

#### Statistical analysis

Unpaired t-tests and one-way ANOVA were performed within each group to determine statistical significance, with an alpha value of 0.05. All statistical analyses were carried out using Prism Version 9 Software (GraphPad).

#### Data availability

The authors confirm that all other data supporting the findings of this study are available within the paper and its supplementary information files.

## Acknowledgments

We thank Ulandt Kim from the NextGen core at Massachusetts General Hospital for the help with single-cell sequencing. We thank Carolyn Smith for outstanding technical support of the immunohistochemistry studies. We thank Dr. Mario Suvá of Massachusetts General Hospital and Dr. Carrie Shawber of Columbia University for important discussions.

## Author contributions

Research design: T.P.P., P.L., K.J.R., K.R., E.R.P. Research performance: P.L., K.J.R., J.J.R, K.R., M.J.O, E.M.B., M.M., A.S.K., G.A.A., M.S.R., H.Z., L.M., M.D., D.J. Data analysis: P.L., K.J.R., M.J.O, E.M.B., A.S.K., G.A.A. Supervision: T.P.P. Writing – original draft: P.L. Writing – review & editing: P.L., K.J.R., J.J.R, M.J.O, A.S.K., L.M., H.K., J.W.B, J.M.U., L.L.M., D.J., T.P.P.

