## Supplemental figures for "Lymphatic muscle cells are unique cells that undergo aging induced changes"

**A** **Figure S1**

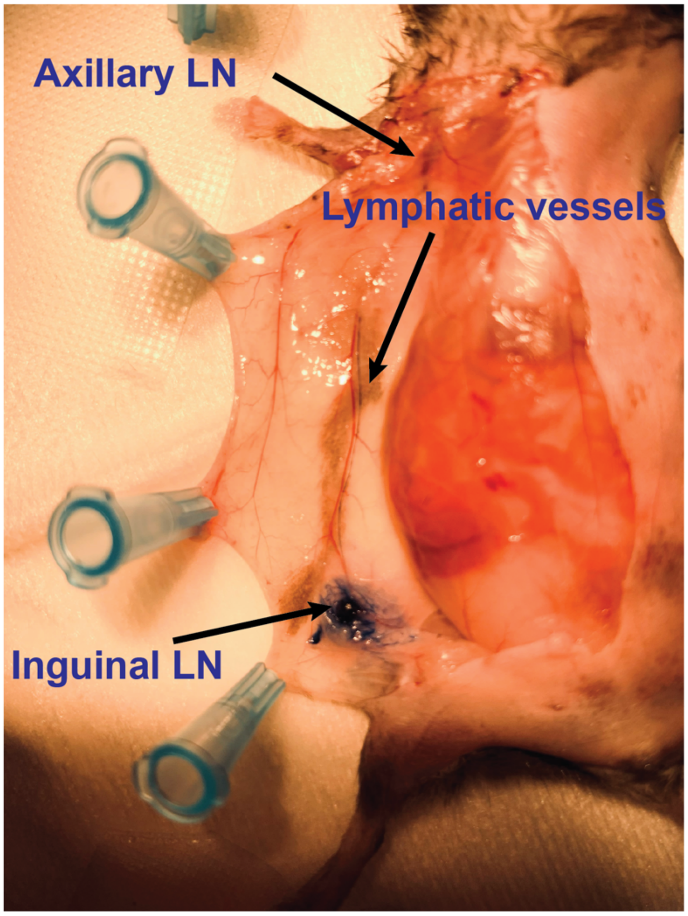


**Figure S1. Identifying the mouse collecting lymphatic vessels.**

**A)** A representative image shows the Evans Blue injection to identify the collecting lymphatic vessel between the inguinal and axillary lymph nodes.

### Figure S2

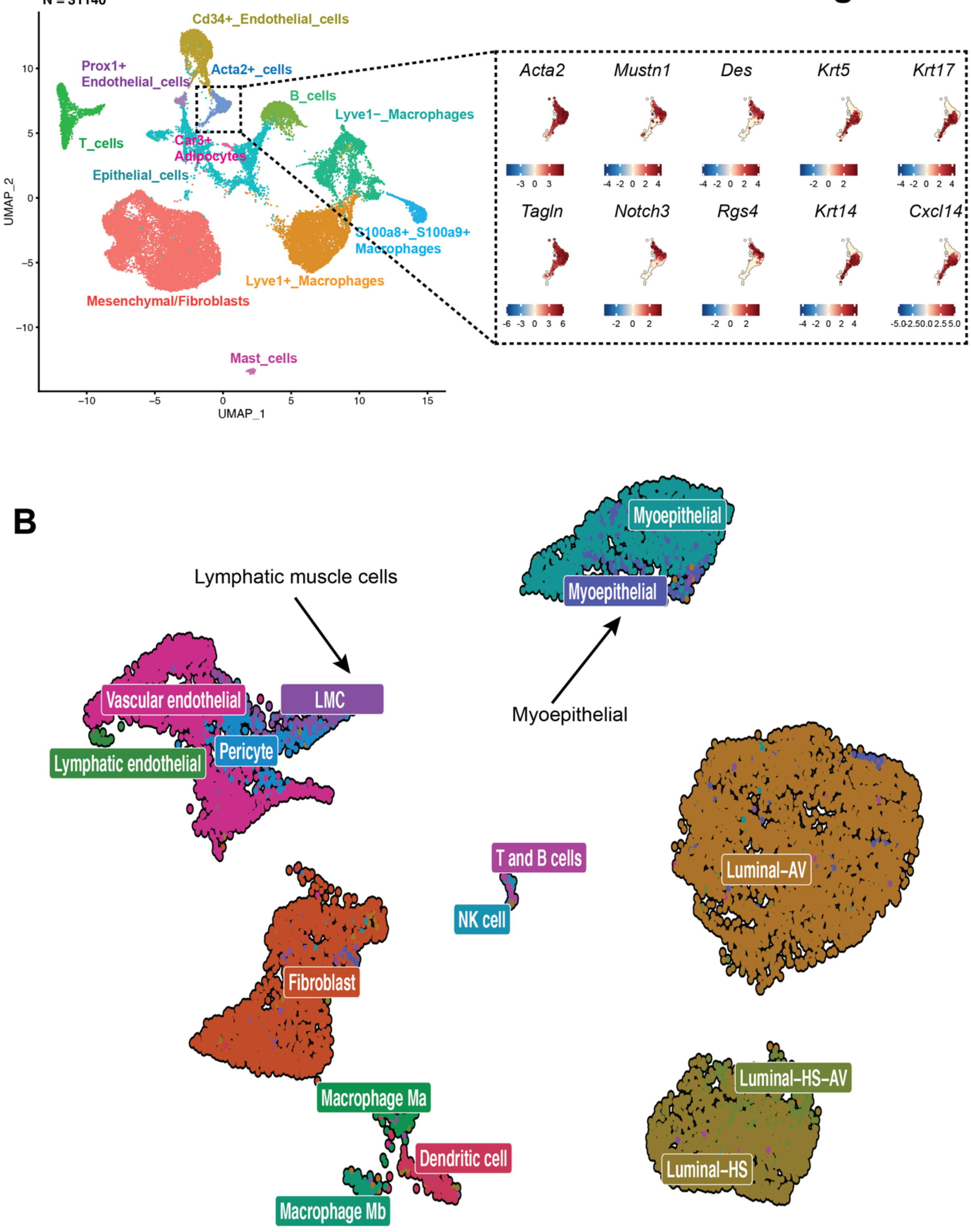
A
N = 31140
Cd34+_Endothelial_cells
Prox1+
Endothelial_cells
Acta2+_cells
B_cells
Lyve1−_Macrophages
T_cells
Car3+
Adipocytes
Epithelial_cells
S100a8+_S100a9+
Macrophages
Lyve1+_Macrophages
Mesenchymal/Fibroblasts
Mast_cells
UMAP_1
UMAP_2
Acta2
Mustn1
Des
Krt5
Krt17
Tagln
Notch3
Rgs4
Krt14
Cxcl14
B
Lymphatic muscle cells
Myoepithelial
Myoepithelial
Myoepithelial
Vascular endothelial
LMC
Pericyte
Lymphatic endothelial
T and B cells
NK cell
Luminal-AV
Fibroblast
Macrophage Ma
Dendritic cell
Macrophage Mb
Luminal-HS-AV
Luminal-HS

**Figure S2. Identifying LMC in murine collecting lymphatic vessel single-cell dataset.**

**A**) UMAP of gene expression of selected marker genes in all *Acta2*+ cells in the collecting lymphatic vessels. Left) The UMAP of all the cells in collecting lymphatic vessels. Right) Single-cell gene expression of selected marker genes was projected into the UMAP of all *Acta2*+ cells. **B)** The UMAP of aggregated *Acta2+* cells from our dataset using lymphatic vessels and cells from mouse mammary fat pad (GSE150580) (Li et al., 2020).

### Figure S3

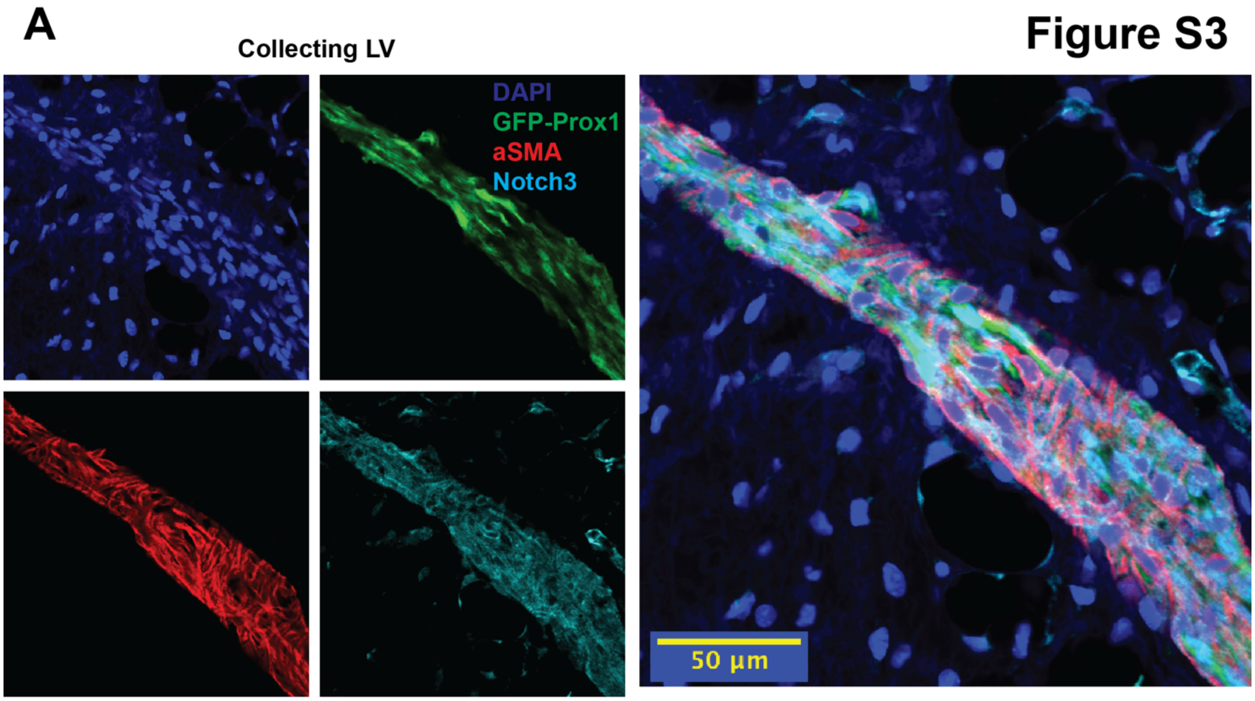


**Figure S3. The presence of Notch3 in LMCs.**

**A)** Representative images of αSMA and Notch3 expression in LMCs of a collecting lymphatic vessel from a Prox1-GFP mice.

**Figure S4**

**A**

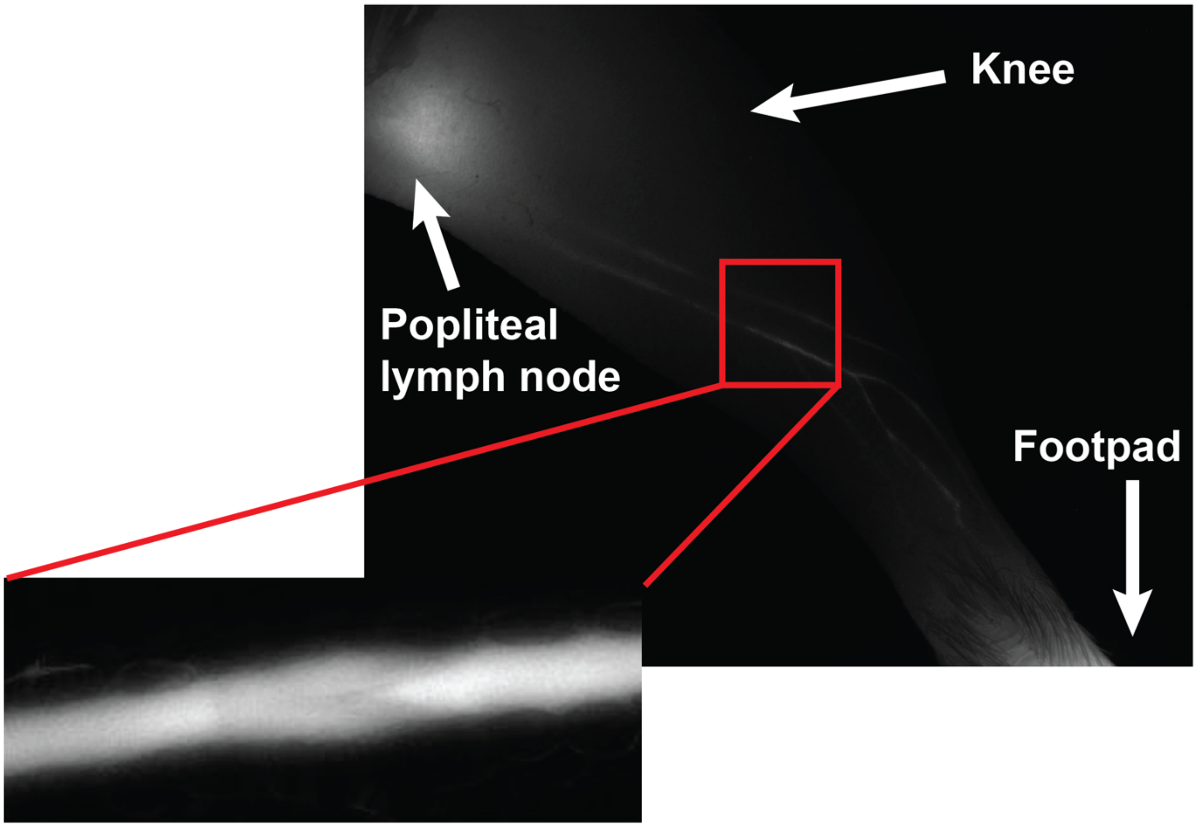
Knee
Popliteal
lymph node
Footpad

**B**

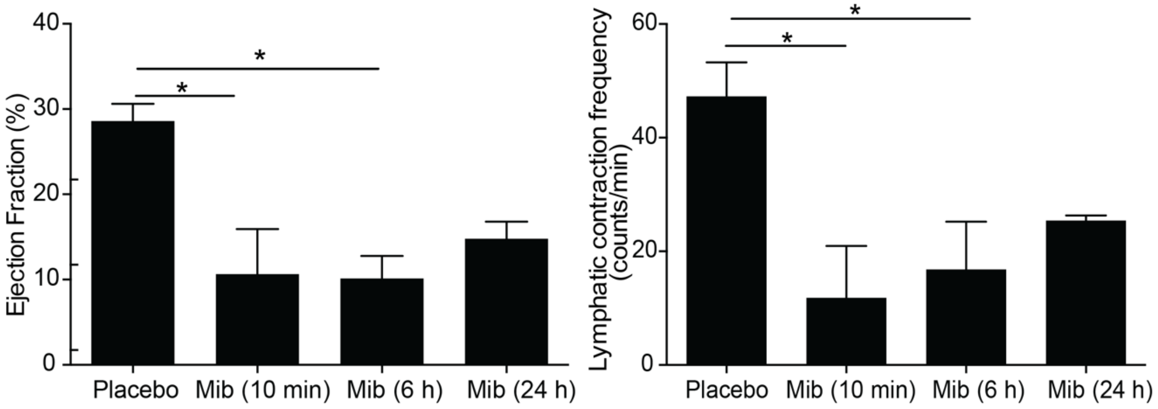
Ejection Fraction (%)
40
30
20
10
0
*
*
Placebo
Mib (10 min)
Mib (6 h)
Mib (24 h)
Lymphatic contraction frequency
(counts/min)
60
40
20
0
*
*
Placebo
Mib (10 min)
Mib (6 h)
Mib (24 h)

**Figure S4. Intravital imaging of lymphatic pumping.**

**A)** Representative images of the intravital imaging of lymphatic vessel pumping in the collecting lymphatic vessels afferent to the popliteal lymph node. **B)** Mibefradil sustains the reduction in ejection fraction and contraction frequency for at least 6 hours.

**Figure S5**

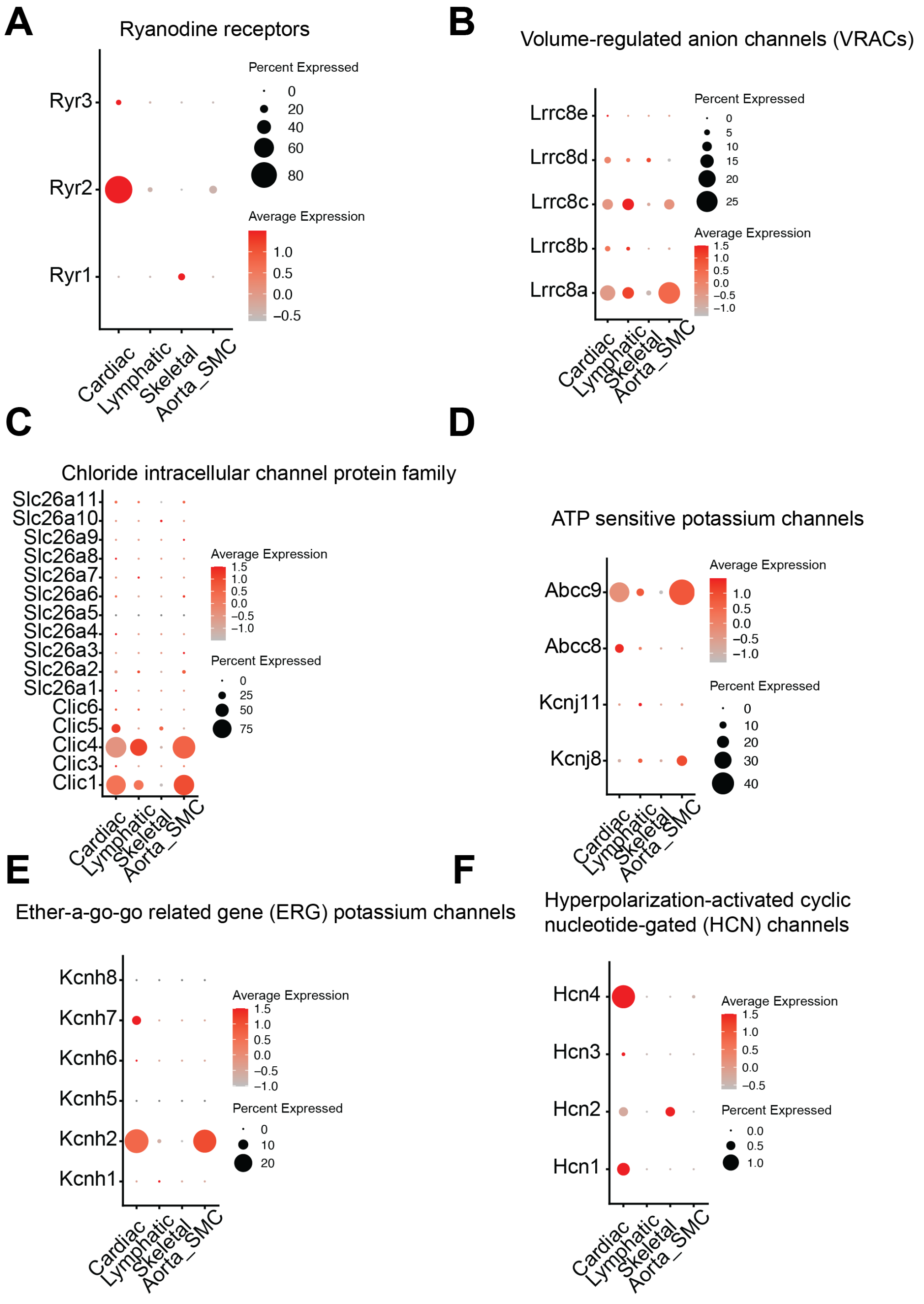


**Figure S5. The presence of anion, chloride, potassium and HCN channel molecules in LMCs.**

**A-F)** The dot plot shows the gene expression of A) Ryanodine receptors, B) voltage-regulated anion channels, C) chloride-intracellular channels, D) ATP-sensitive potassium channels, E) Ether a-go-go related gene (ERG) potassium channels and F) HCN channels in all the muscle cells.

**Figure S6**

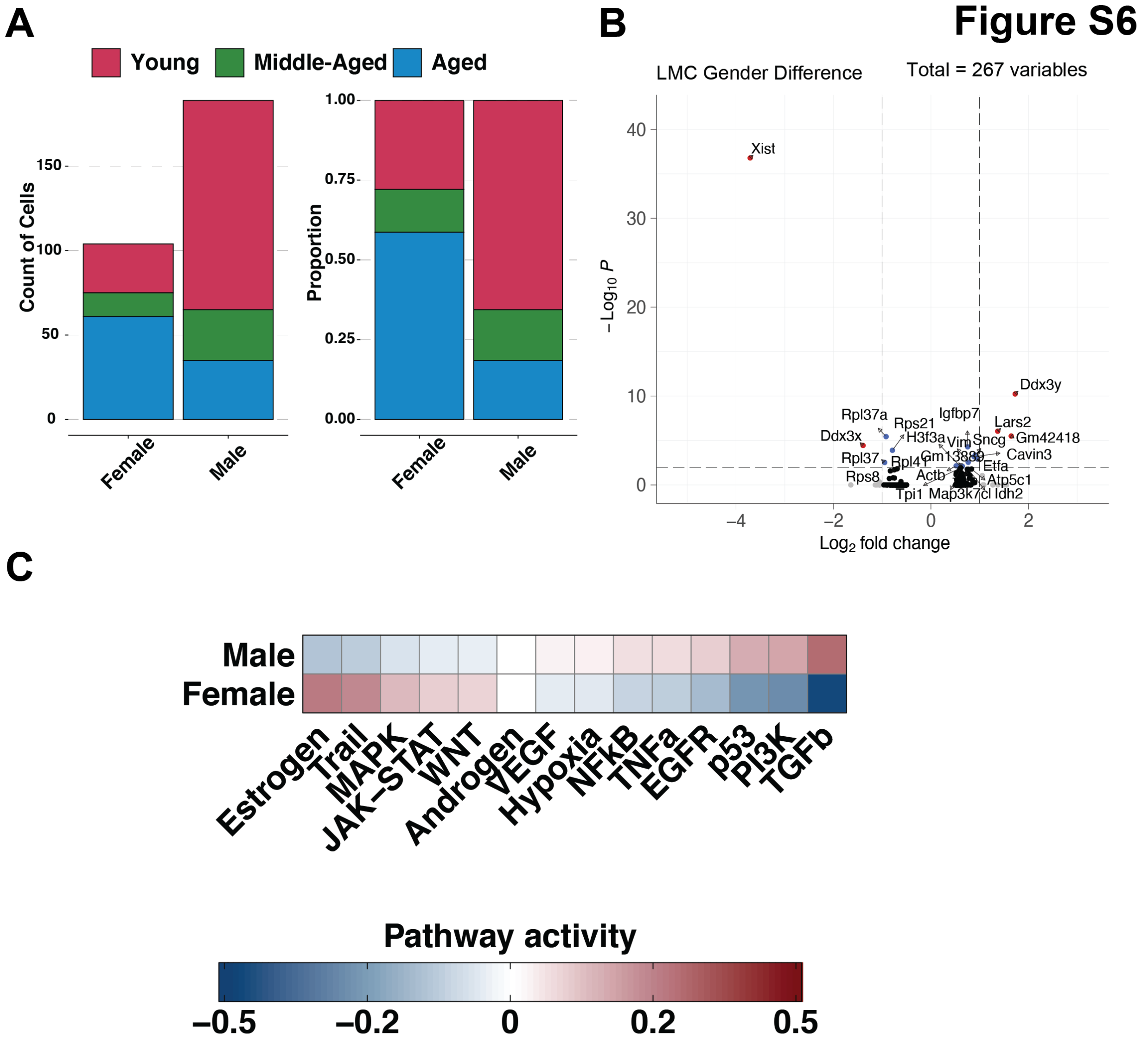


**Figure S6. The difference between LMCs in male and female mice.**

**A)** The composition of cells in LMCs grouped by sex. Left) The number of cells annotated by age and sex. Right) The proportion of cells grouped by sex. **B)** The volcano plot depicts the differentially expressed genes in LMCs between male and female mice. **C**) The PROGENy pathway analysis of the differentially expressed genes in LMCs between male and female mice.

A

### Figure S7

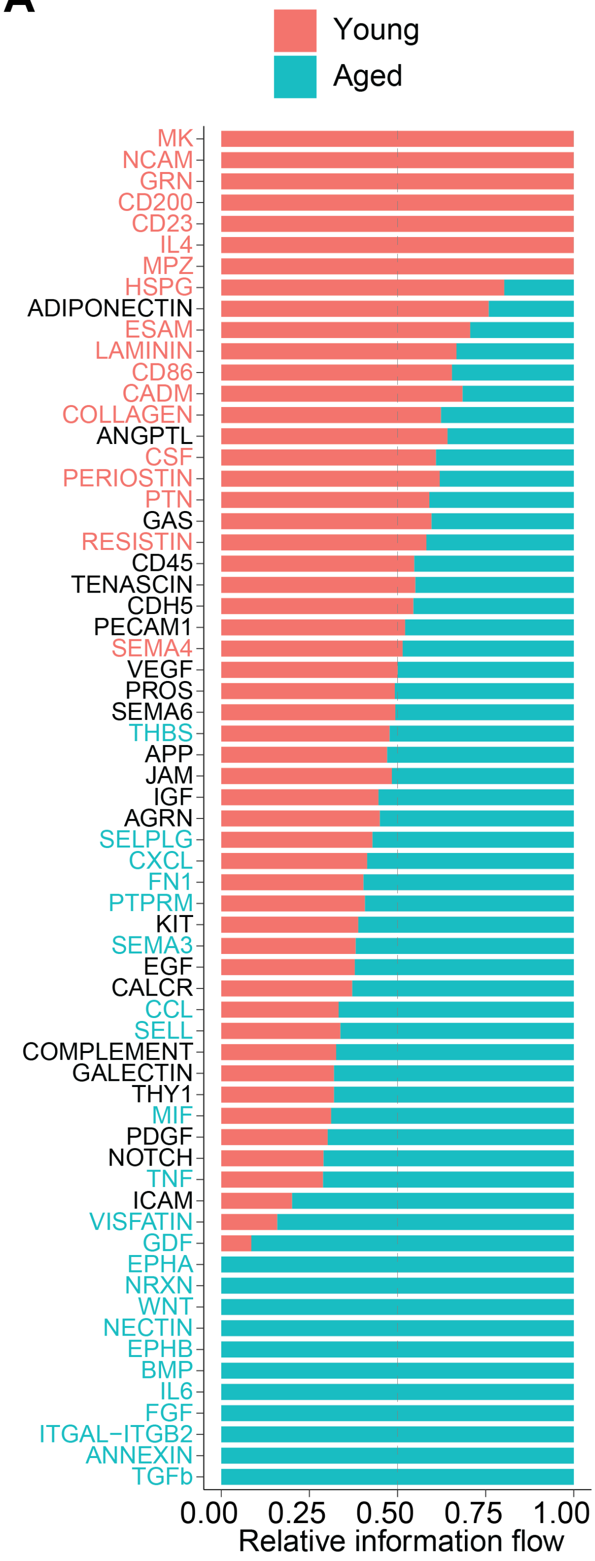

Young
Aged
MK
NCAM
GRN
CD200
CD23
IL4
MPZ
HSPG
ADIPONECTIN
ESAM
LAMININ
CD86
CADM
COLLAGEN
ANGPTL
CSF
PERIOSTIN
PTN
GAS
RESISTIN
CD45
TENASCIN
CDH5
PECAM1
SEMA4
VEGF
PROS
SEMA6
THBS
APP
JAM
IGF
AGRN
SELPLG
CXCL
FN1
PTPRM
KIT
SEMA3
EGF
CALCR
CCL
SELL
COMPLEMENT
GALECTIN
THY1
MIF
PDGF
NOTCH
TNF
ICAM
VISFATIN
GDF
EPHA
NRXN
WNT
NECTIN
EPHB
BMP
IL6
FGF
ITGAL−ITGB2
ANNEXIN
TGFb
0.00
0.25
0.50
0.75
1.00
Relative information flow

**Figure S7. The cell-cell communication signaling in lymphatic vessels.**

**A)** The stack bar plot represents the sum of communication probability among all pairs of cell groups in the young (6-week-old) and aged (18-month-old) groups. The significant signaling pathways are ranked based on differences in the overall cell-cell interaction strength in the inferred networks between young (6-week-old) and aged (18-month-old) lymphatic vessels. The top signaling pathways colored by red are enriched in the young (6-week-old) group, while those colored by green are enriched in the aged (18-month-old) group.
