## Supplemental Tables for "Lymphatic muscle cells are unique cells that undergo aging induced changes"

**Table 1. The quality control of single-cell sequencing datasets.**

| Sample | Group | Total Reads | Number of Cells | Number of Genes | Average Reads per Cell | Average Genes per Cell | Average Percent Mitochondrial RNA |
|---|---|---|---|---|---|---|---|
| LV1 | Young (6-week-old male) | 93,543,578 | 9,785 | 16,455 | 2,705 | 1,119 | 4.10 |
| LV2 | Young (6-week-old female) | 74,925,516 | 5,970 | 16,562 | 2,627 | 1,087 | 3.39 |
| LV3 | Middle-Aged (12-month-old male) | 84,391,934 | 3,639 | 15,731 | 3,677 | 1,376 | 2.79 |
| LV4 | Middle-Aged (12-month-old female) | 88,490,555 | 1,864 | 16,128 | 6,398 | 1,630 | 3.43 |
| LV5 | Aged (18-month-old male) | 99,271,958 | 1,207 | 14,735 | 6,667 | 1,834 | 3.38 |
| LV6 | Aged (18-month-old female) | 75,981,072 | 8,675 | 16,824 | 2,377 | 1,008 | 4.91 |

**Table 2. The summary of the number of cells in each sample.**

| | **Young (6-week-old female)** | **Young (6-week-old male)** | **Middle-Aged (12-month-old female)** | **Middle-Aged (12-month-old male)** | **Aged (18-month-old female)** | **Aged (18-month-old male)** |
|---|---|---|---|---|---|---|
| **Mesenchymal/Fibroblasts** | 1951 | 4685 | 492 | 1795 | 3259 | 496 |
| **Lyve1+_Macrophages** | 965 | 1701 | 323 | 628 | 1905 | 104 |
| **Epithelial_cells** | 315 | 487 | 353 | 157 | 1625 | 79 |
| **Lyve1-_Macrophages** | 480 | 1105 | 251 | 199 | 746 | 37 |
| **Cd34+_Endothelial_cells** | 260 | 1096 | 65 | 144 | 193 | 130 |
| **T_cells** | 636 | 272 | 171 | 226 | 378 | 27 |
| **B_cells** | 1160 | 108 | 38 | 67 | 103 | 20 |
| **S100a8+_S100a9+_Macrophages** | 50 | 66 | 71 | 262 | 111 | 176 |
| **Myoepithelial** | 53 | 27 | 62 | 0 | 227 | 3 |
| **Lymphatic_muscle_cells (LMCs)** | 29 | 124 | 14 | 30 | 61 | 35 |
| **Prox1+_Endothelial_cells (LECs)** | 25 | 22 | 7 | 88 | 21 | 73 |
| **Mast_cells** | 33 | 74 | 5 | 15 | 43 | 5 |
| **Adipocytes** | 13 | 18 | 12 | 28 | 3 | 22 |
